# High-resolution small RNAs landscape provides insights into alkane adaptation in the marine alkane-degrader *Alcanivorax dieselolei* B-5

**DOI:** 10.1101/2022.11.09.515887

**Authors:** Guangshan Wei, Sujie Li, Sida Ye, Zining Wang, Kourosh Zarringhalam, Jianguo He, Wanpeng Wang, Zongze Shao

## Abstract

Alkanes are widespread in the ocean, and *Alcanivorax* is one of the most ubiquitous alkane-degrading bacteria in the marine ecosystem. Small RNAs (sRNAs) are usually at the heart of regulatory pathways, but sRNA-mediated alkane metabolic adaptability still remains largely unknown due to the difficulties of identification. Here, differential RNA sequencing (dRNA-seq) modified with a size selection (∼50-nt to 500-nt) strategy was used to generate high-resolution sRNAs profiling in the model species *Alcanivorax dieselolei* B-5 under alkane (*n*-hexadecane) and non-alkane (acetate) conditions. As a result, we identified 549 sRNA candidates at single-nucleotide resolution of 5’-ends, with 63.4% of transcription start sites (TSSs) and 36.6% of processing sites (PSSs). These sRNAs originated from almost any locations in the genome, regardless of intragenic (65.8%), antisense (20.6%) and intergenic (6.2%) regions, and RNase E may function in the maturation of sRNAs. Most sRNAs locally distribute across the 15 reference genomes of *Alcanivorax*, and only 7.5% of sRNAs are broadly conserved in this genus. Expression responses to alkane of several core conserved sRNAs, including 6S RNA, M1 RNA and tmRNA, indicate that they may participate in alkane metabolisms and result in more actively global transcription, RNA processing and stresses mitigation. Two novel CsrA-related sRNAs are identified, which may be involved in the translational activation of alkane metabolism-related genes by sequestering the global repressor CsrA. The relationships of sRNAs with characterized genes of the alkane sensing (*ompS*), chemotaxis (*mcp, cheR, cheW2*), transporting (*ompT1, ompT2, ompT3*) and hydroxylation (*alkB1, alkB2, almA*) were created based on the genome-wide targets prediction. Overall, the sRNAs landscape lays the ground for uncovering cryptic regulations in the critical marine bacterium, among which both core and species-specific sRNAs are implicated in the alkane adaptive metabolisms.

## 1. Introduction

Alkane, as a ‘double-edged sword’, fuels human society but threatens ecological balance. In the ocean, alkanes are ubiquitous and can originate from both natural and anthropogenic activities, such as cyanobacterial biosynthesis (Lea-Smith et al., 2015), geothermal processes (He et al., 2021), seafloor seepages (Scoma et al., 2017), and spills during the oil drilling and transporting (Yakimov et al., 2022). *Alcanivorax*, as one of the most widespread and abundant hydrocarbon-degrading bacteria in the ocean, is particularly good at metabolizing various alkanes, regardless of aliphatic, branched, halo- and cyclo-alkanes (Li and Shao, 2014; Wang and Shao, 2014; Scoma et al., 2017; Gregson et al., 2019; Yakimov et al., 2022). Members of *Alcanivorax* play critical roles in the marine ecosystem through regulating the alkanes cycling.

For decades, a number of studies have focused on the isolation, diversity and distribution of *Alcanivorax* based on the cultivation and culture-independent methods (Yakimov et al., 1998; Liu and Shao, 2005; Liu et al., 2019; Yakimov et al., 2022), and the key enzymes and pathways of alkane metabolism have been uncovered, especially in the *A. borkumensis* and *A. dieselolei* species (Liu et al., 2011; Li and Shao, 2014; Wang and Shao, 2014; Barbato et al., 2016; Gregson et al., 2019). Multiple key enzymes usually coexist in their genomes, such as the AlkB, AlmA and cytochrome P450 alkane hydroxylases, to deal with different alkanes substrates (Liu et al., 2011; Wang and Shao, 2013; Wang and Shao, 2014; Barbato et al., 2016; Gregson et al., 2019; Yakimov et al., 2022). In particular, the key genes in alkane sensing, chemotaxis, signal transduction, transporting and regulation have been revealed, and an entire network of alkane metabolism from sensing to degradation has been constructed in the model alkane-degrader *A. dieselolei* B-5 (Wang and Shao, 2014).

It has been well established that sRNAs are widely distributed both in various gram-positive and gram-negative bacteria, and they play diverse and widespread roles in almost all the cell processes (Carrier et al., 2018; Wassarman, 2018; Boutet et al., 2022; Ponath et al., 2022). Bacterial sRNAs are at the heart of regulatory pathways, which allow bacteria to rapidly acclimate to the ever-changing environments through reshaping the related genes expression and adjusting their metabolism (Förstner et al., 2018; Wassarman, 2018). Generally, sRNAs can regulate gene expression through three different mechanisms: base-pairing to target RNAs (e.g., *trans*- and *cis*-acting sRNAs), directly interacting with proteins (e.g., 6S RNA and CsrA protein related sRNAs), and intrinsic functions (e.g., catalytic sRNAs) (Wassarman, 2018; Krieger et al., 2022). Given that the highly dynamic environments and fluctuated alkanes content in the ocean (Lea-Smith et al., 2015; Scoma et al., 2017), transient regulations via sRNAs are supposed to be essential to the quick adaptation for the ubiquitous and predominant alkane-degraders. However, knowledge of sRNA-mediated alkane metabolic regulations remains quite limited.

Bacterial sRNAs are usually regarded as the ‘dark matter’ in genome, due to their enormous diversity but difficult to identify. Firstly, most sRNAs are poorly conserved except those within phylogenetically close species (Lindgreen et al. 2014), resulting in the bad performance of bioinformatic predictions to most cases (Barik and Das, 2018; Boutet et al., 2022). Secondly, the classical mutation-based genetic screening strategy to investigate protein-coding genes is usually inefficient for detecting sRNAs due to lacking of discernible phenotypes for most disrupted sRNAs (Sridhar and Gunasekaran, 2013; Rachwalski et al., 2022). Importantly, more and more evidences support that sRNAs can originate from any locations in the genome. In addition to the well-known intergenic and antisense regions, myriad sRNAs are derived from the UTRs (untranslated regions) and CDSs (coding sequences) of mRNAs (Miyakoshi et al., 2015; Dar and Sorek, 2018; Adams and Storz, 2020; Hoyos et al., 2020; Tang et al., 2020; Adams et al., 2021; Krieger et al., 2022; Ponath et al., 2022). Fortunately, with the rapid development of high-throughput sequencing technologies, RNA-seq-based approaches have been created to detect the global sRNA profiling (Hör et al., 2018), especially the differential RNA sequencing (dRNA-seq), which can even capture the sRNAs overlapped with other transcripts (Yu et al., 2018).

The dRNA-seq is an emerging technology for global mapping of transcription boundaries and transcript origins with single-nucleotide resolution (Sharma et al., 2010; Sharma and Vogel, 2014; Bischler et al., 2015). Owing to the unique advantage of differentiating transcription start site (TSS) and RNA processing sites (PSS), dRNA-seq could detect sRNAs originated from any locations in the genome (Yu et al., 2018). Here, to identify the sRNA candidates in oil-degrading bacteria and infer the roles of sRNAs in alkane metabolisms, dRNA-seq was used to capture the sRNA landscape in a model hydrocarbonclastic bacterium *A. dieselolei* B-5. Furthermore, we combined gene expression responses to alkane and targets analysis, to predict the sRNA candidates and corresponding regulatory mechanisms in alkane metabolisms. The results provide insights into sRNA-mediated alkane adaptability of the prevalent marine alkane-degrader.

## 2. Results and Discussion

### 2.1. dRNA-seq uncovering genome-wide sRNA profiles with high-resolution of 5’-ends

First of all, to verify the accuracy of 5’-end of transcripts in dRNA-seq, we compared the start sites of highly conserved tRNAs predicted by two independent methods (tRNAscan-SE 2.0 and ARAGORN) with the results annotated by dRNA-seq (Laslett and Canback, 2004; Schwengers et al., 2021) (Table S1). As a result, the vast majority of tRNAs (42/49) were predicted exactly at the same start sites by the two methods, in which most of them (39/42) have a precisely mapped TSS or PSS in the dRNA-seq results (Table S1). Not only the relative small tRNAs (with sizes of ∼70-nt to 100-nt), but also the larger conserved tm-RNA (358-nt) predicted by the Bakta showed the same start site with the dRNA-seq results (Table S1). Other few unmapped tRNAs might be caused by the incorrect prediction or silenced expression in the detected conditions. Moreover, the relative expression levels of the matched tRNAs and tm-RNA spanned a great range of six orders of magnitude (with TPM from 1 to 10^6^), indicating that both rare and abundant transcripts are detected.

Therefore, the dRNA-seq could capture the 5’-end of transcripts with different sizes and varied expression levels at the single-nucleotide resolution in this study. To gain comprehensive profiles of the sRNAs, we combined the ANNOgesic prediction and manual inspection based on the dRNA-seq and ssRNA-seq data (Yu et al., 2018; Ponath et al., 2021). Finally, a total of 549 sRNA candidates were obtained in strain B-5 under both alkane and non-alkane conditions (Figure 1A and Table S2). Given the accurate and unique 5’-end locations on the genome, the sRNAs are named in the form of sRNAXXX, of which XXX stands for the corresponding specific genomic location of each sRNA (Table S2). Generally, most of the sRNAs (∼85%) ranged from 100-nt to 300-nt in length, with a median of 205-nt (Figure 1B). For the strand-distribution, sRNAs showed more minus-strand preference than CDSs in the genome (Figure 1C). Among these sRNAs, 2.6% of them (14/549) showed high similarities (>70%) with at least one other sRNA detected in this study, indicating that few sRNAs redundantly expressed in B-5 genome. Interestingly, nearly half of these high similarity homologs (6/14) are as anisense transcripts of different transposase genes (Table S3). This suggests that the transposition activities of transposons may be widely repressed at post-transcriptional level through antisense sRNAs in B-5 genome, which is also reported in other bacterial and archaeal species before (Ellis and Haniford, 2016).

**Figure 1.**
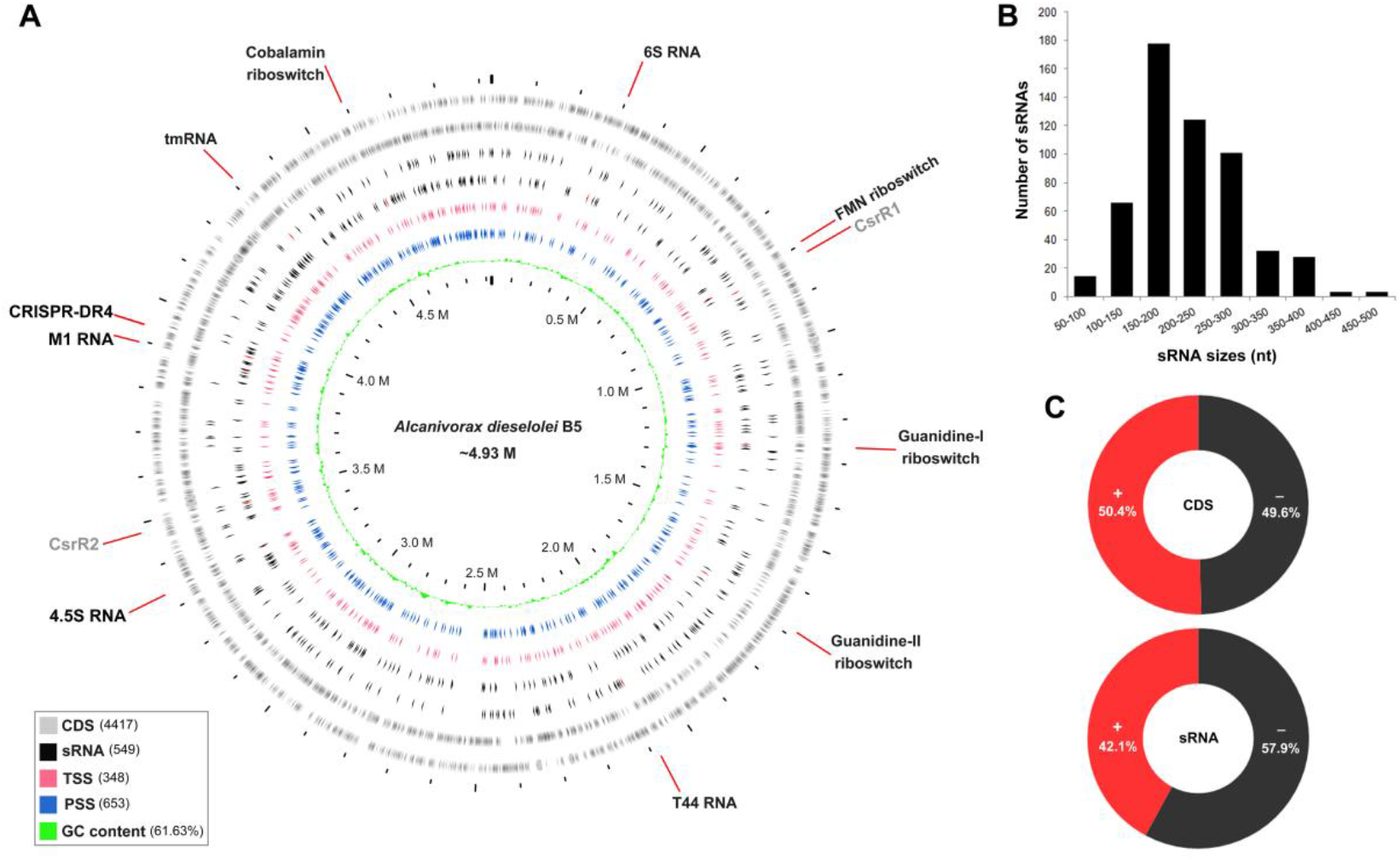
Genome-wide sRNAs landscape of *A. dieselolei* B-5. (**A**) Diagram shows the distributions of sRNAs, TSSs and PSSs across the B-5 whole genome. From outside to inside, the rings represent the distributions of CDSs, sRNAs, TSSs, PSSs and the GC content respectively, and the values in brackets show corresponding numbers or pencentage in the genome. The orientations of each ring are clockwise for the plus (+) strand and counterclockwise for the minus (−) strand. The relatively conserved sRNAs are red marked on the corresponding rings, and their annotated names are displayed on the outermost of the plot, black fonts for Rfam families and grey for identified CsrA-related sRNAs. Circos plot was created by using Proksee (https://proksee.ca/). (**B**) The size distribution of sRNAs. (**C**) Comparison of the distributing proportions of CDSs and sRNAs in two strands (+/-) of the genome.

### 2.2. Origin patterns of sRNAs associated with promoters and RNase E cleavages

To understand the origin patterns of sRNAs in the genome, considering the accurate TSSs and PSSs obtained by dRNA-seq, we first classified the sRNAs based on their 5’-end origins into two types: tssRNA (transcription <u>s</u>tarting <u>sRNA</u>) and pssRNA (<u>p</u>roce<u>s</u>sing <u>sRNA</u>) (Figure 2A and Table S4). The tssRNAs originated from their own transcription start sites account for approximately two-thirds of the total sRNAs, which are more abundant than pssRNAs derived from the processing of other transcripts (Figure 2A). For the 348 TSSs, sequence logo of the upstream 50-nt displayed typical motif characteristics of bacterial promoters in the -10 and -35 regions (Figure 2B). Meanwhile, iPromoter-2L-based prediction indicated that about half of the upstream regions of TSSs could be recognized by distinct sigma factors, especially by σ^70^ (40.2%) and σ^24^ (12.4%) (Table S5). Furthermore, an obvious A/G purine preference at the TSS (+1 position) was also detected (Figure 2B), which is well consistent with previous reports from other bacterial transcription start sites (Feklıstov et al., 2014; Liu et al., 2018).

**Figure 2.**
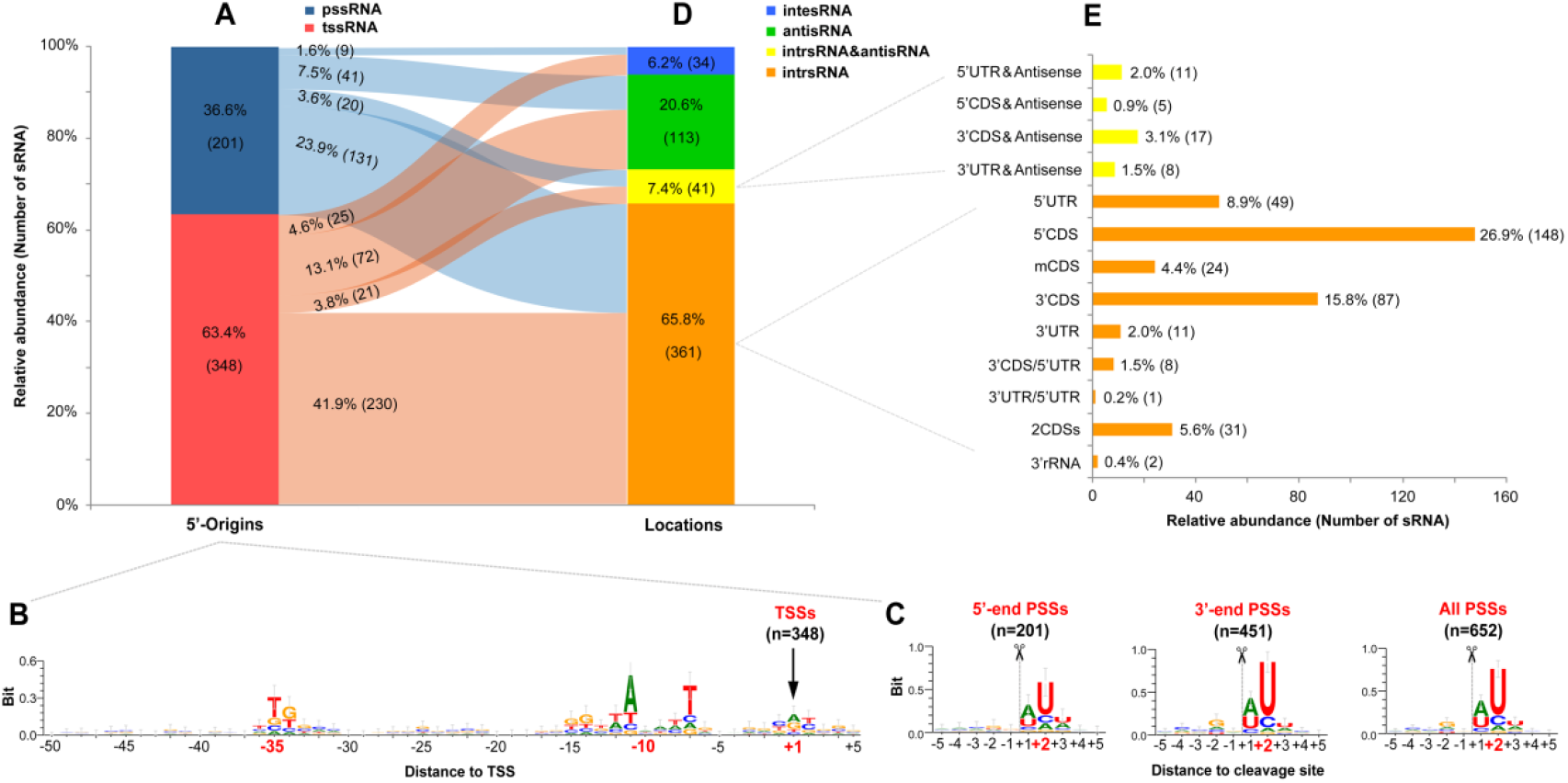
Origin and location patterns of sRNAs. (**A**) Distribution of sRNAs classified by the origins of 5’-ends. (**B**) and (**C**) show the sequence logos of neighboring regions of TSSs and PSSs, respectively. The “-” and “+” before the numbers represent the upstream and downstream positions separately, and the featured positions are shown with bold and red fonts. (**D**) and (**E**) show the distributions of sRNAs classified by their locations relative to the genomic annotations. The percentage represents the ratio of one type of sRNA in the total, and the following number in the bracket shows the corresponding amount. The relationships between (A) and (D) are connected by a Sankey diagram created by using the SankeyMATIC (https://sankeymatic.com/build/).

For the PSSs at the 5’- and 3’-ends of sRNAs, the sequence logos of their neighboring sequences showed a strong preference for uridine (U) at the downstream 2-nt (+2 position) of cleavage sites, and a 5-nt consensus motif of “RN↓WUU” (with “↓” for “cleavage site”, “R” for “G/A”, “W” for “A/U”, and “N” for “any nucleotide”) was identified (Figure 2C and Table S6), reminiscent of the essential RNase E in Gram-negative bacteria which prefers to cleave RNA in single-stranded A/U-rich regions (Mackie, 2013; Mohanty and Kushner, 2018). Unexpectedly, the motif feature of *A. dieselolei* matches the RNase E-specific recognition motifs in diverse bacteria very well (Figure S1), including *Salmonella enterica* (Chao et al., 2017), *Vibrio cholerae* (Hoyos et al., 2020) and even the model cyanobacterium *Synechocystis* sp. PCC 6803 (Hoffmann et al., 2021). This highlighted that RNase E should be the overriding player in the processing of sRNAs in *A. dieselolei*, and the inference is further supported by the highly conserved crucial residues for specific recognition and cleavage of RNase E between *A. dieselolei* and the aforementioned bacteria (Figure S1). These results indicate that the origin patterns of tssRNAs and pssRNAs are closely associated with the upstream promoters and RNase E cleavages, respectively.

### 2.3. Diverse location patterns relative to the annotated genes in B-5 genome

To understand the sources of the sRNAs in B-5 genome, they were also classified based on their locations relative to the annotated genes. As a result, the sRNAs distribute in diverse genomic locations and can be divided into three major types: intesRNA (<u>inte</u>rgenic <u>sRNA</u>), antisRNA (<u>anti</u>sense <u>sRNA</u>) and intrsRNA (<u>intr</u>agenic <u>sRNA</u>) (Figure 2D and Table S4). The intesRNA is located on the intergenic region of two nearby genes and without overlap with transcripts of the two genes; the antisRNA overlaps with transcript of annotated gene but with divergent transcriptional direction; the intrsRNA overlaps with partial transcript of annotated gene and with same transcriptional direction (See detailed examples in Figure S2). According to the specific location of overlaps, the intrsRNAs can be further subdivided into sRNAs in 5’UTR (at the 5’UTR of mRNA), 5’CDS (overlapped with the 5’-end of CDS), mCDS (at the middle of CDS), 3’CDS (overlapped with the 3’-end of CDS), 3’UTR (at the 3’UTR of mRNA), 3’rRNA (overlapped with the 3’-end of rRNA), 2CDSs (overlapped with two same directional CDSs) and so on (Figures 2E, S2 and Table S4). In addition, some other sRNAs that overlap with two transcripts of nearby genes in divergent directions are designated as intrsRNA&antisRNA (Figures 2D, 2E and S2). The majority of our retrieved sRNAs belonged to intrsRNAs (65.8%), particularly sRNAs overlapped with the 5’- and 3’-ends of mRNAs (Figures 2D and 2E), indicating that mRNA may be the primary origination source of sRNAs in *A. dieselolei*. This finding is in congruence with recent evidences in different bacteria, such as sRNAs from 3’-UTRs in *Salmonella enterica* (Chao et al., 2017), 5’UTRs and CDSs in *Escherichia coli* (Dar and Sorek, 2018; Adams et al., 2021), and CDSs, 5’UTRs as well as 3’-UTRs in *Xanthomonas campestris* (Tang et al., 2020).

### 2.4. Locations instead of origin patterns associated with sRNA features

The sRNA features, including sequence length, GC content and minimum free energy were calculated and compared according to the above classifications. The sequence lengths of all sRNAs ranged from 61-nt to 470-nt, with an average of 217±72 -nt (Figure 3A and Table S7). For different types of sRNAs, tssRNA (217±72 -nt) and pssRNA (217±7 1-nt) showed no significantly difference in length (Wilcoxon rank sum test, P > 0.05), while the antisRNAs (244±81 -nt) and intesRNAs (193±84 -nt) had notably longer and shorter sizes than others, respectively (P < 0.05, Table S7). In a comparative investigation based on 816 identified sRNAs across 33 bacterial species, the authors summarized the average lengths of *cis-* and *trans*-sRNAs are 219- and 182-nt, respectively (Barik and Das, 2018), which is very similar to the length of the distributions of corresponding antisRNA and intesRNA in our study.

**Figure 3.**
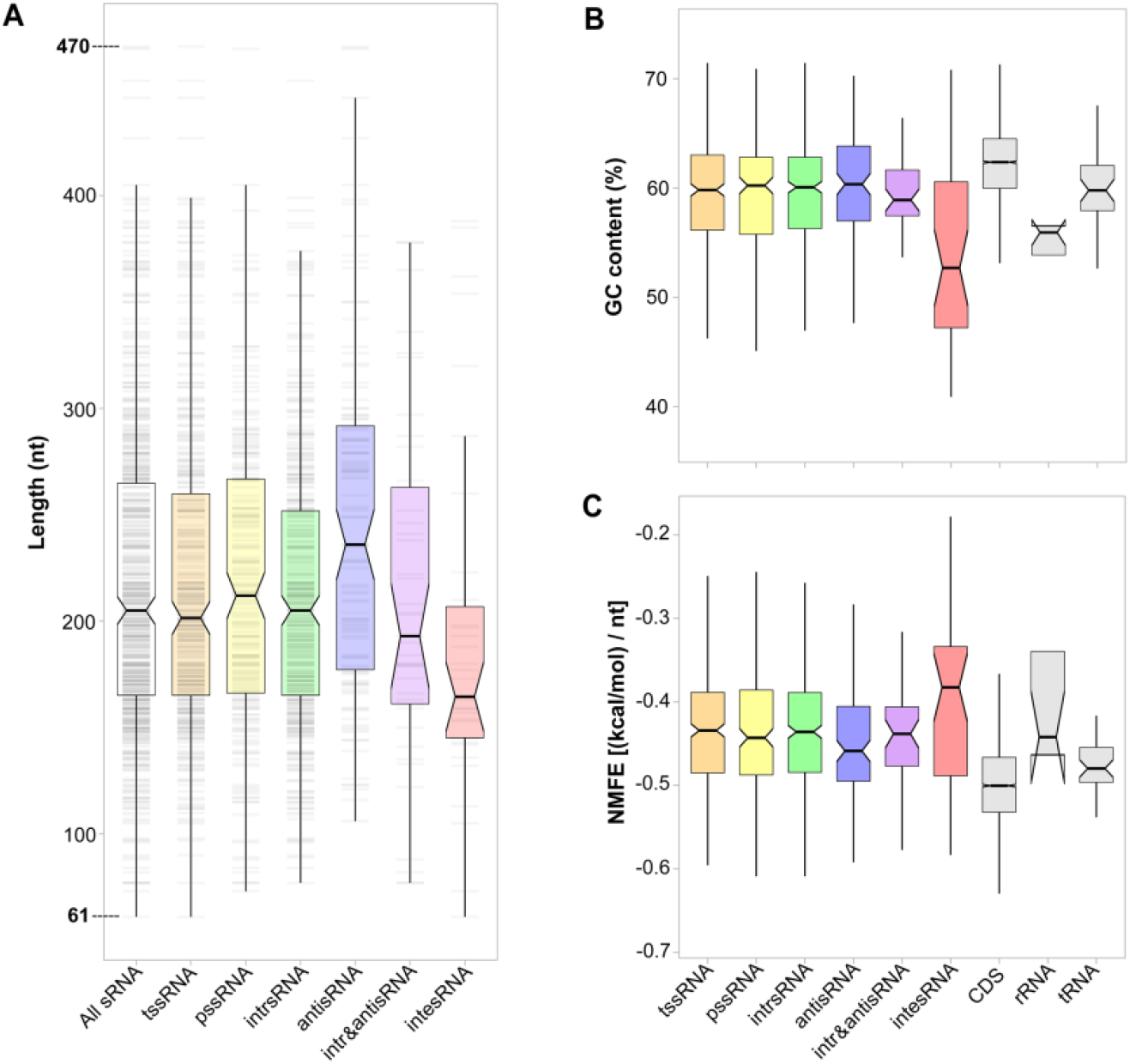
Sequence and structure features of sRNAs with different classifications. (**A**) Boxplot shows the length distributions of different types of sRNAs. The grey stripes in the background show the overall length distributions. (**B**) GC contents and (**C**) NMFEs of classified sRNAs, CDSs, rRNAs and tRNAs.

The intesRNAs have the lowest GC content (53.9% for average), which is comparable to rRNAs (55.4%) but significantly lower than other types of RNAs (58.4-61.8%). Except the intesRNA, all the other types of sRNAs had similar GC contents (58.4%-59.8%), which are significantly lower than CDSs (61.8%), higher than rRNAs (55.4%) and comparable with tRNAs (60.1%) (Figure 3B and Table S7). However, both integenic and antisense sRNAs usually had relatively lower GC contents than the CDSs, rRNAs and tRNAs in other bacterial species (Barik and Das, 2018; Ryan et al., 2020). Of note, most of the previously reported sRNAs are usually derived from the genomes of human gut bacteria, such as *Escherichia, Salmonella*, and *Bacteroides*, which have considerably lower GC contents (∼54% for average) than our marine derived strain B-5 (61.6%) (Barik and Das, 2018; Ryan et al., 2020). Although the GC content has been used as predictive information for intergenic sRNA identification in some bacteria (Livny and Waldor, 2007), our results indicated that the GC contents of sRNAs vary in different genomic locations and distinct species.

The secondary structure feature of sRNA is reflected by the length normalized minimum free energy (NMFE), which is closely correlated with the structural folding stability of RNA (Barik and Das, 2018). Except between intesRNA (−0.3993 (kcal/mol) / nt for average) and antisRNA (−0.4478), all the different types of sRNAs showed no significantly difference in NMFEs. However, all the sRNAs showed remarkably higher NMFEs (ranging from -0.3993 to -0.4478) than that of the CDSs (−0.4955) and tRNAs (−0.4800), and were comparable with the rRNAs (−0.4148) (Figure 3C and Table S7). The higher NMFEs of sRNAs may correlate with their less stable secondary structures, since the sRNA-chaperone Hfq protein is also a key factor correlated with their stability through protecting sRNAs from degradation (Wagner, 2013; Barik and Das, 2018; Bar et al., 2021). Furthermore, the NMFEs of different types of RNAs showed a strongly negative correlation with their average GC contents (Spearman’s ρ=-0.95, P<0.001) in this study (Table S7), indicating the close relationship between RNA structure stability and GC contents.

Taken together, the sequence- and structure-based features of sRNAs are less influenced by their origin patterns (i.e., transcription starting or processing origins) but are closely correlated with their location patterns (i.e., intergenic, intragenic or antisense) in *A. dieselolei*. It is noted that these features may provide some valuable parameters for the bioinfomatic prediction of sRNAs in genome, the differences caused by the species-dependent specificity should be fully considered.

### 2.5. Protein-coding potential of sRNAs

GeneMarkS-2 prediction indicated that more than half of the sRNAs (302/549) have potential to encode protein-residues (without start and/or stop codons), further Pfam annotations revealed that these residues are mainly due to the overlaped CDSs of intrsRNAs (Table S8). Ribo-seq result showed that only 6.9% sRNAs (38/549) contain typical small ORFs (sORFs) (Table S9), and this proportion is similar with a previous report on several bacterial phyla (5-15%) by a machine learning prediction method (Friedman et al., 2017). BLASTP results further showed that most of the coding sRNAs were hypothetical proteins or without known homolog in the NCBI nr database (Table S9). Intriguingly, additional 139 sRNAs even without coding potential were also detected in the Ribo-seq data (Table S9). Similar phenomenon was also observed in the Ribo-seq results of *E. coli*, and the persistent existence of sRNAs was attributed to their protective secondary structures instead of the ribosome binding (Fremin and Bhatt, 2020). Therefore, both the bioinformatic prediction and the Ribo-seq results support that most of the sRNAs are still non-coding in this study. Considering the difficulty of sORF validation (Friedman et al., 2017), the limitation of Ribo-seq (Fremin and Bhatt, 2020), and the bifunctional (both sORF-coding and base-pairing regualtion) potential (Carrier et al., 2018), these sRNAs with coding potential were not excluded from the next regulatory analyses.

### 2.6. Identification of Rfam-annotated core sRNAs

To reveal the sRNAs with homologs in the database of known RNA families, all the sRNA candidates were aligned in the Rfam database. As a result, 11 of the 549 sRNAs were successfully annotated into 10 Rfam families, including a 6S RNA, a 4.5S RNA, a M1 RNA, a tmRNA, a T44 RNA, two CRISPR RNA direct repeat elements (CRISPR-DR4) and four different riboswitches (Figures 1A, 4A and Table S10). All these sRNAs were detected in both the dRNA-seq and ssRNA-seq results with clear read peaks (Figure 4A), and they are mainly originated from the intergenic or 5’-UTR regions with their own TSSs (Table S10). BLASTN similarity searches against the 15 reference genomes of *Alcanivorax* suggested that only four of them, i.e., 6S RNA, 4.5S RNA, M1 RNA and tmRNA, are widely distributed in *Alcanivorax* genus (Figure 4B). In fact, these four sRNAs are also broadly existed in other bacteria and play key roles as core sRNAs, and their detailed features in *Alcanivorax* are as follows.

**Figure 4.**
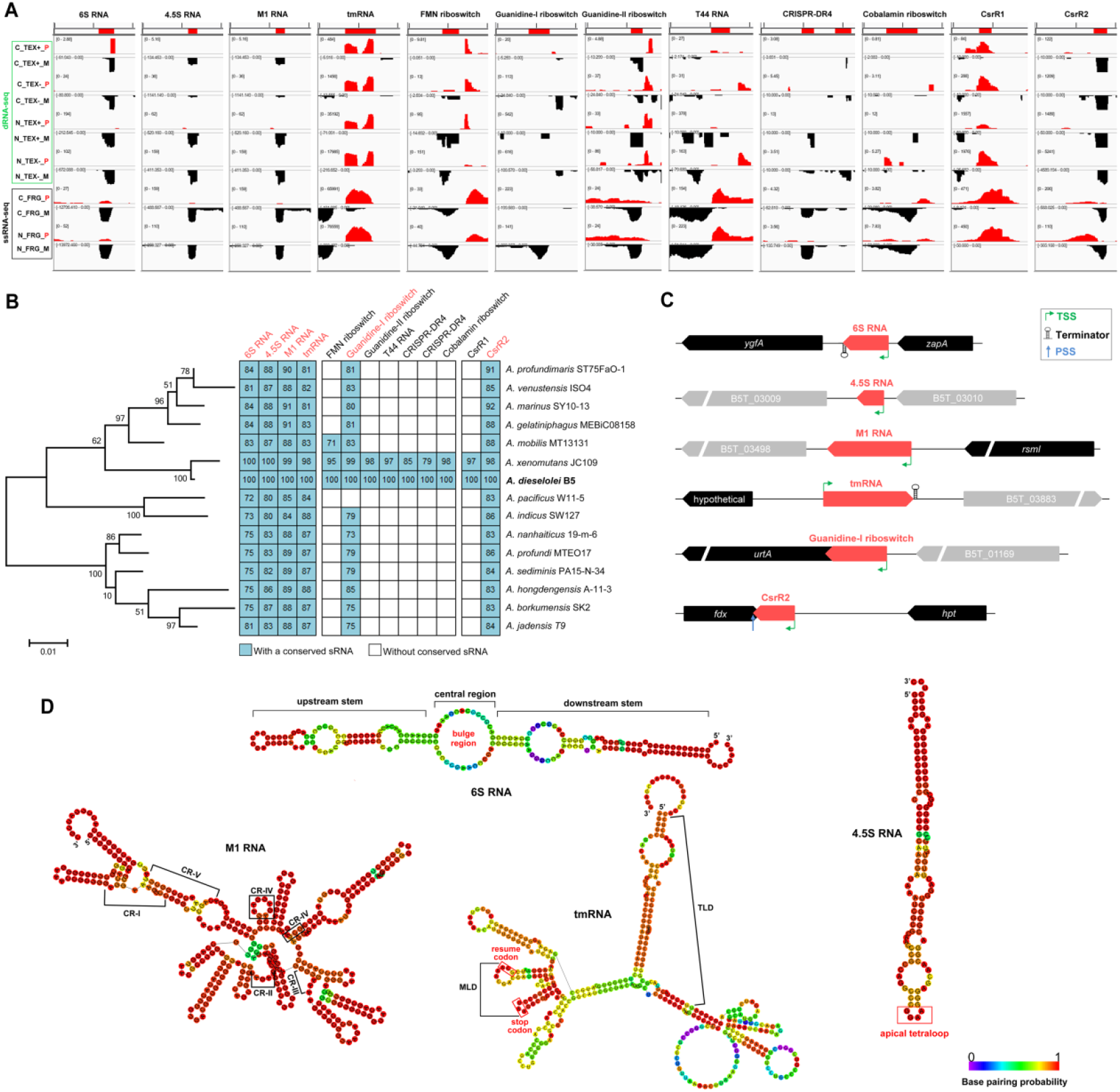
Rfam-annotated sRNAs of *A. dieselolei* and their distributions in genus *Alcanivorax*. (**A**) IGV views on the read coverages of Rfam-annotated sRNAs. Tracks display the different experimental conditions (i.e., C for C16 (hexadecane) and N for NaAc (sodium acetate) carbon sources) and RNA-seq strategies (i.e., TEX+ and TEX-for dRNA-seq and FRG for ssRNA-seq). The reads mapped on the plus (P) and minus (M) strands of genome are in red and black colours, respectively. (**B**) The homologs of Rfam-annotated sRNAs in Alcanivorax. Left part shows the 16S rRNA gene-based maximum-likelihood phylogenic tree of representative Alcanivorax species constructed using MEGA6 (www.megasoftware.net), the bootstrap values that are more than 50 are presented on the related branches. Middle part shows the distributions and identities (%) of sRNAs in different species, and the names of broadly conserved sRNAs are in red. Right part shows the names of corresponding species. (**C**) Neighboring gene synteny of highly conserved sRNAs. The red arrows in the middle represent the sRNAs, black for the highly conserved nearby genes and grey for the less conserved genes. The arrow direction to left indicates genes on the minus-strand and to the right for the plus-strand. The details of the nearby genes are listed in Table S13. (**D**) Secondary structures and conserved features of the core sRNAs.

#### 2.6.1. 6S RNA and associated pRNAs

The 6S RNA is an important global riboregulator at the transcriptional level through mimicing the open DNA promoter to sequester the RNA polymerase (RNAP) in *Escherichia coli* and *Bacillus subtilis* (Wassarman, 2018). Since lacking of conserved primary sequence, it is usually hard to predict the 6S RNA gene in other non-model bacteria (Ponath et al., 2021). Here, combined dRNA-seq and Rfam alignment, a 197-nt 6S RNA was first identified from *A. dieselolei* and is conserved in *Alcanivorax* with varied sequence similarities (Figure 4B). Gene synteny and Rfam annotation of all the homologs further confirmed the conservation of 6S RNA in *Alcanivorax* (Figure 4C and Table S11). The upstream neighboring gene is *zapA* encoding cell division protein, and the downstream gene *ygfA* encoding 5-formyltetrahydrofolate cyclo-ligase that is involved in purine metabolism (Figure 4C and Table S12). The *zapA-6S RNA*-*ygfA* synteny is also reported in other bacterial taxa, especially the *Gammaproteobacteria* (Wehner et al., 2014), implying some underlying relationships between 6S RNA and cell division and nucleic acid metabolism. Further secondary structure prediction indicated that *A. dieselolei* 6S RNA displays a structure of “upstream stem - central bulge - downstream stem” (Figure 4D), which is in line with the typical structure of *E. coli* 6S RNA (Chen et al., 2017). Importantly, we obtained evidence for the existence of 6S RNA-associated product RNAs (pRNAs) nearby the central bulge region (Figure S3), which may mediate the release of sequestered RNAP by 6S RNA (Wassarman, 2018). In fact, four putative pRNAs with different 5’-ends and varied lengths were uncovered in *A. dieselolei* (Figure S3), which are more diverse than previously described, like in *Helicobacter pylori* and *Bacteroides thetaiotaomicron* (two potential pRNAs), in *E. coli* and *B. subtilis* (one pRNA), and in *Legionella pneumophila* (without pRNA) (Sharma et al., 2010; Wassarman, 2018; Ryan et al., 2020). The diverse pRNAs may contribute to the flexible release of the sequestered RNAP by 6S RNA to rapidly recover the transcriptional activity globally, which is critical to the survival of *Alcanivorax* in the ever-changing marine environments.

#### 2.6.2. 4.5S RNA of SRP

The bacterial signal recognition particle (SRP) RNA, termed 4.5S RNA in *E. coli*, is the core component of SRP for delivering inner membrane proteins to the translocon or the insertase for membrane insertion (Steinberg et al., 2018; Ponath et al., 2021). The 113-nt 4.5S RNA of *A. dieselolei* showed 80-88% sequence identities with that in most of the *Alcanivorax* species (Figure 4B), and all the homologs were confirmed in the Rfam database (Table S11). However, the neighboring gene synteny of 4.5S RNA is not conserved among *Alcanivorax* species (Figure 4C and Table S12). The secondary structure of 4.5S RNA forms a hairpin-like structure with several internal loops and a conserved apical GGAA tetraloop (Figure 4D), well resembles that of *E. coli* (Steinberg et al., 2018).

#### 2.6.3. M1 RNA of RNase P

M1 RNA is the core catalytic component of RNase P in bacteria that participates in the processing and maturation of various RNAs, like tRNAs, tmRNA, 4.5S RNA, and some mRNAs (Bechhofer and Deutscher, 2019; Gößringer et al., 2021; Mohanty and Kushner, 2022). We obtained a 362-nt M1 RNA from *A. dieselolei*, which has widespread homologs with high similarities in *Alcanivorax* type strains (Figure 4B and Table S11). A highly conserved *rsmI* gene encoding 16S rRNA (cytidine (1402)-2’-O)-methyltransferase is located at the upstream of M1 RNA gene, but the downstream genes are not conserved in *Alcanivorax* (Figure 4C and Table S12). Moreover, the five universally <u>c</u>onserved regions (CR-I to CR-V) and key catalytic sites of M1 RNA were also identified in *Alcanivorax* (Figures 4D and S4), further confirming its functional conservation as the core component of RNase P resembling other identified bacteria (Mondragón, 2013 ; Gößringer et al., 2021).

#### 2.6.4. tmRNA and its flexible MLD

Another core sRNA from *A. dieselolei* is a 388-nt transfer-messenger RNA (tmRNA), which plays a vital role in rescuing the stalled ribosomes on defective mRNAs in bacteria (Guyomar et al., 2021). The *Alcanivorax* tmRNAs shared a conserved upstream gene encoding a hypothetical protein in an opposite transcriptional direction (Figures 4C and S5). Like in *E. coli*, a key mRNA-like domain (MLD) was also identified in *A. dieselolei* tmRNA (Figure 4D), which features an internal ORF encoding a small protein (10 amino acids) to tag the problematic peptides produced by the stalled ribosomes for proteases recognition (Guyomar et al., 2021). Interestingly, the resume codon (GCA) and stop codon (UAA) of MLD are highly conserved in *E. coli* and *Alcanivorax* species (Figure 4D), but the fourth and the fifth codons encoded different amino acids with similar physicochemical properties (Glu-Asn for *E. coli* and Asp-Thr/Ser for *Alcanivorax*) (Figure S5). Moreover, nearly half of the MLD codons of *Alcanivorax* showed base wobbles mainly in the third position, but most of the corresponding amino acids are still unchanged (Figure S5). Different bacterial species use distinct codons to encode flexible protein-tags in MLD of tmRNA probably depends on their codon preferences, and the functions of the protein-tags may retain constant due to the degeneracy of codon and similar properties of amino acids.

### 2.7. Identification of Rfam-annotated local sRNAs

Except the above four core sRNAs, some Rfam-annotated local sRNAs that are conserved in partial bacteria are also identified from *A. dieselolei*.

#### 2.7.1. Riboswitches related to detoxication of guanidine and metabolism of B vitamins

Riboswitch is one of the most common riboregulators in bacteria, which recognizes specific small molecular ligand by its aptamer domain and modulates the downstream gene transcription or translation through switching the conformation (Pavlova et al., 2019). Guanidine-I riboswitch is widely distributed in most species of *Alcanivorax* and conservatively located at the upstream of an *urtA* gene encoding a putative urea ABC transporter substrate-binding protein (Figures 4B and 4C). While the Guanidine-II riboswitch is only detected in *A. dieselolei* and its closest species *A. xenomutans*, with a downstream gene encoding a SugE-like protein (Figure 4B and Table S10). Guanidine is derived from the metabolisms of arginine, creatine or guanine, which is a toxic compound to the cell (Nelson et al., 2017). Guanidine riboswitches play crutial roles in degradation or export this toxin through regulating expressions of the downstream guanidine carboxylase or transporter genes (Nelson et al., 2017; Battaglia and Ke, 2018). Considering the downstream genes and distribution patterns, most *Alcanivorax* members may be capable of exporting the intracellular guanidine via the sensing and regulation of Guanidine-I riboswitch, and *A. dieselolei* and *A. xenomutans* may have more powerful guanidyl-scavenging ability due to their extra Guanidine-II riboswitch.

The other two riboswtiches are closely related to the metabolism of B vitamins, with FMN riboswitch and Cobalamin riboswitch for the regulation of biosynthesis or transport of riboflavin (vitamin B_2_) and cobalamin (vitamin B_12_), respectively (Polaski et al., 2016; Wilt et al., 2020). Although Cobalamin riboswitch and FMN riboswitch are among the top-three-most-widely distributed riboswitches in bacteria (Polaski et al., 2016; Pavlova et al., 2019), they showed limited distribution in *Alcanivorax* (Figure 4B). Based on the functions of downstream genes (Table S10), we deduce that *A. dieselolei* and the closely relatives may modulate their intracellular vitamin B_2_ and B_12_ concentrations through FMN- and Cobalamin-riboswitches mediated riboflavin biosynthesis and cobalamin uptake, respectively. Note that despite most of the known riboswitches control gene expression through a *cis*-acting way (Pavlova et al., 2019), the above four riboswitches in *Alcanivorax* showed clearly accumulated transcription peaks (Figure 2A), and two of them have PSSs at their 3’-ends (Table S10), indicating their *trans*-acting regulation potential as sRNAs, which is also reported in other bacteria (Vogel et al., 2003; Loh et al., 2009; Ponath et al., 2021).

#### 2.7.2. T44 RNA likely associated with factors of translation process

T44 RNA is first reported in *E. coli* in a microarray-based transcriptome study about twenty years ago (Tjaden et al., 2002), and it was found widely distributed in different bacteria but with unknown function ever since (Toffano-Nioche et al., 2012). Here we also identified a T44 RNA from *A. dieselolei*, locating between the genes *map* (encoding methionine aminopeptidase involved in the amino-terminal maturation of translation process) and *rpsB* (encoding 30S ribosomal protein S2) (Figure 4A and Table S10), which is consistent with most bacteria (Toffano-Nioche et al., 2012). Intriguingly, based on the dRNA-seq results, T44 RNA is antisense and overlapped to the 5’-UTRs of *map* and *rpsB* genes, respectively (Figure S6). Meanwhile, no more TSS, but a few PSSs were detected in the downstream of T44 RNA, suggesting that T44 RNA gene together with the two downstream *rpsB* and *tsf* (encoding translation elongation factor Ts) genes may form an operon (Figure S6), which is similar to the *Vibrio splendidus* (Toffano-Nioche et al., 2012). Given that all the related *rpsB, tsf* and *map* genes are involved in the translation processes, we propose that T44 RNA may influence the global translation through *cis*-acting effects on these genes in *A. dieselolei*.

#### 2.7.3. Two active CRISPR RNAs within a Type I-F CRISPR-Cas system

In *A. dieselolei*, we also identified two species-specific CRISPR RNAs (crRNAs) of CRISPR-DR4 family, which is characterized by the same repeat with palindrome to form a 6-bp stem-loop with high GC contents (Figures 4A, 4B and S7). CRISPR-Cas system is the adaptive immune system in most prokaryotes against the phages as well as mobile genetic elements (Hille et al., 2018). Based on the CRISPRCasFinder (Couvin et al., 2018), a Type I-F CRISPR-Cas system was identified in *A. dieselolei* genome, and both detected crRNAs are within a large CRISPR array containing 63 distinct spacers (Figure S7). The 159-nt crRNA sRNA3935639 is originated from a TSS and contains three different spacers separated by two repeats, while the 209-nt sRNA3936114 is derived from a PSS and has four spacers separated by three repeats (Table S10 and Figure S7). Compared to the common mature crRNA that usually has one spacer-repeat pair, here the two crRNAs with multiple spacer-repeat pairs indicate a complex processing mechanism of CRISPR array with larger stable intermediates, which is also seen in type I-B systems of bacterial *Fusobacterium nucleatum, Clostridium thermocellum* and archaeal *Methanococcus maripaludis* (Richter et al., 2012; Ponath et al., 2021). Interestingly, both the active crRNAs contain one spacer that has a putative seed sequence targeting the integrated phage genes with 14-nt perfect matches in *A. dieselolei* genome (Figure S7), and both the targeted genes belong to tailed and double-stranded DNA bacteriophages of order *Caudovirales* (Table S13).

### 2.8. Distribution characteristics of sRNA homologs across Alcanivorax species

Except a few Rfam-annotated sRNAs, most of the sRNAs (538/549) detected in *A. dieselolei* are truly novel and without known homologs in the Rfam database. To further clarify the homologous distributions of all the detected sRNAs, BLASTN analysis was carried out with a threshold of 75% sequence identity across the 15 representative species of genus *Alcanivorax*. As a result, most of the sRNAs are highly specific to *A. dieselolei* and its close species, with more than 80% sRNAs distributing in less than half of (≤7) the *Alcanivorax* species, especially in *A. dieselolei* and *A. xenomutans* (∼60%) (Figure 5A and Table S14). Notably, a total of 41 sRNAs (7.5%) are broadly conserved among diverse *Alcanivorax* species with 75-100% sequence identities (Figures 5A and 5B). Interestingly, most of the broadly conserved sRNAs’ identies showed significantly positive correlations (Spearman-based analyses, P < 0.05 or 0.01) with their 16S rRNA-based phylogenies (Table S15). The intesRNAs showed a higher proportion in the broadly conserved sRNAs (12.2%, 5/41) than that in the whole sRNAs (6.2%, 34/549) (Figures 2 and 5B). Except the Rfam-annotated core sRNAs, an antisense transcript of tmRNA (sRNA4307718) was also detected (Figure 5B), indicating an undetermined regulation role relevant to the tmRNA. In addition, an intesRNA (sRNA4758272) located between genes encoding ribosome biogenesis GTP-binding protein (B5T_04289) and Na^+^/H^+^ antiporter subunit G (B5T_04290) is also highly conserved in *Alcanivorax*, and even with two homologous copies in some species (Figure 5B). In fact, this phenomenon of multiple copies was also found in some other broadly and poorly conserved sRNAs (Figure 5B and Table S14), suggesting these sRNAs may play indispensable roles in their host *Alcanivorax* species.

**Figure 5.**
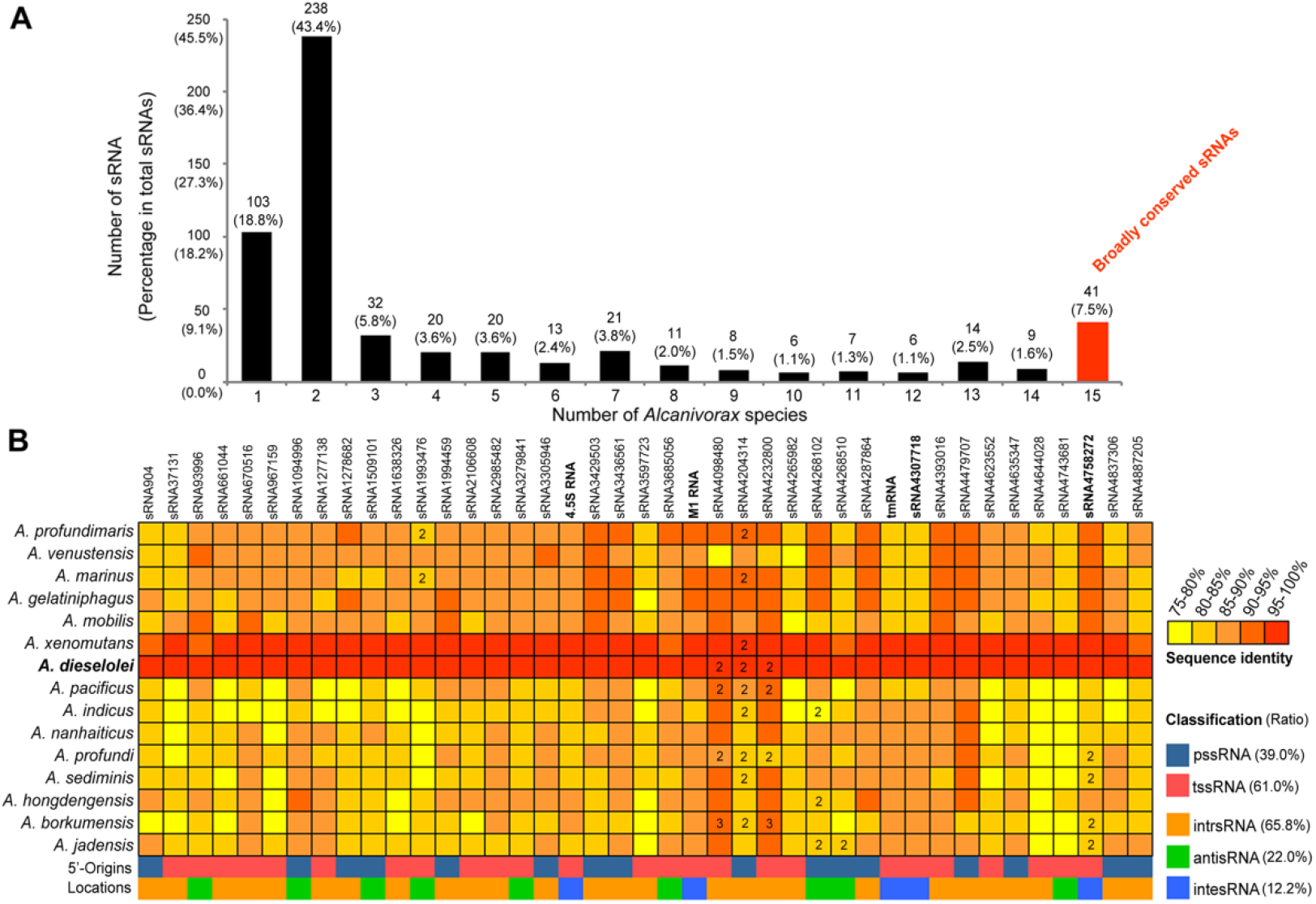
Distribution of sRNA homologs across different species of *Alcanivorax*. (**A**) The number (percentage) distribution of sRNA homologs across one (*A. dieselolei*-specific) to all 15 (broadly conserved) species of *Alcanivorax*. The bar with broadly conserved sRNAs is marked in red. (**B**) The sequence identity distribution of the 41 conserved sRNAs across different Alcanivorax species and their classifications in the reference strain B-5. The different sequence identities are shown using color gradients from yellow to red. The number (2 or 3) in the grid represents the copy number of related sRNA in corresponding species, and the average identity of multiple copies is shown. The classifications of sRNAs are also displayed by distinct colors.

### 2.9. Active expressions of sRNAs facilitating A. dieselolei adapt to alkane utilization

Overall, the expression levels of the 549 sRNAs showed very broad ranges, spanning 6-7 orders of magnitudes under the alkane and non-alkane conditions (Table S16). The top 50 highly expressed sRNAs of each condition represented approximate 90-99% of the total number of hits in all samples (Table S16), and nearly half of them (22/50) are overlapped in both conditions with or without hexadecane as growth substrate (Figure 6).

**Figure 6.**
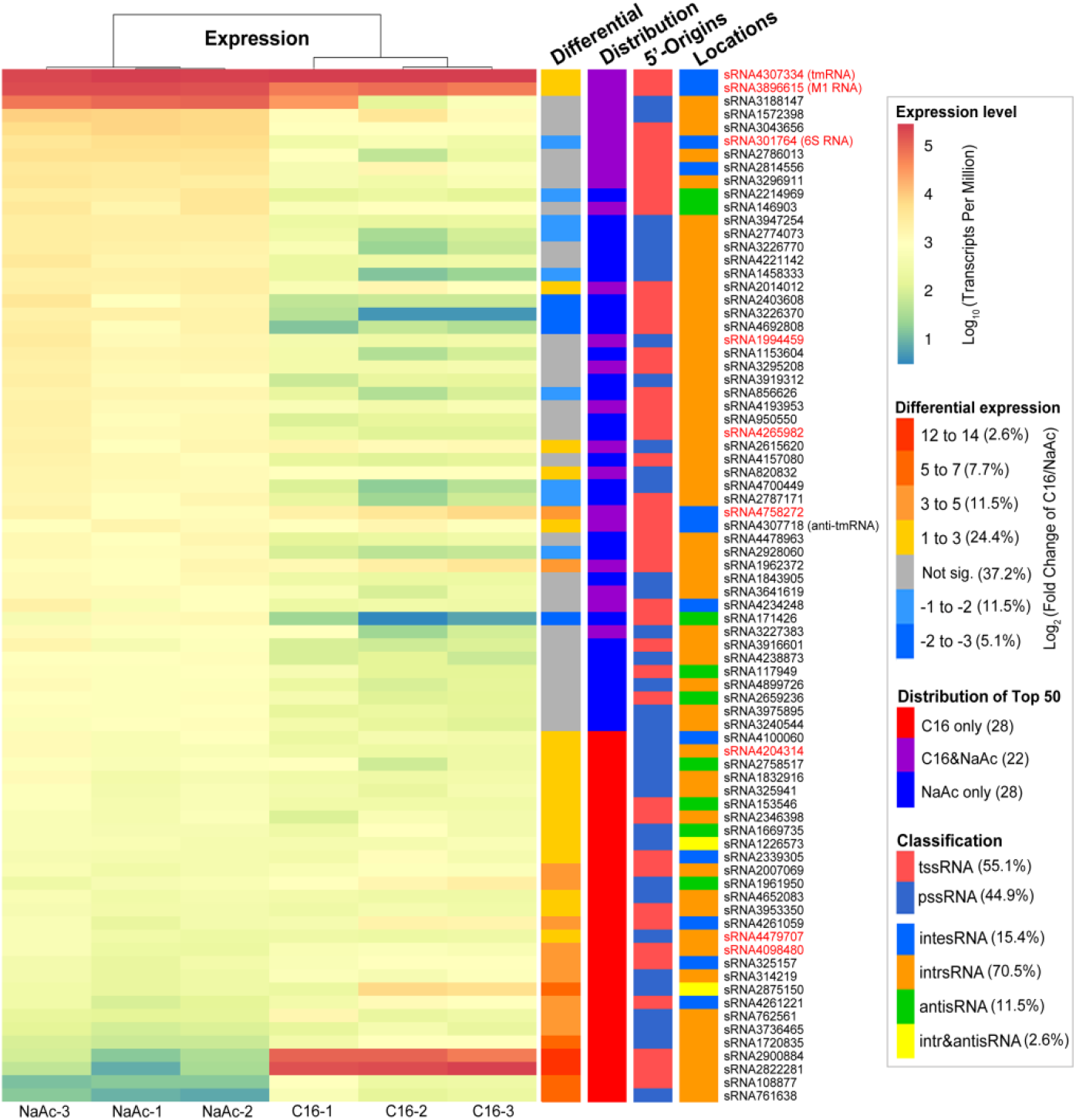
Expression of the top 50 sRNAs in alkane and non-alkane conditions. From left to right, the heatmap and colored bar charts show the expression levels in each sample of NaAc (non-alkane) or C16 (alkane) as carbon source, the differential expression of alkane vs. non-alkane conditions, the distribution of the top 50 highly expressed sRNAs in the two conditions and the classifications based on sRNA origins and locations. The sRNA names are listed behind, and the broadly conserved ones are in red.

More than half (54.8%) of the 549 sRNAs showed similar expression levels, irrespective of alkane or acetate as the sole carbon and energy source, while 127 sRNAs (23.1%) were notably up-regulated and 121 (22.1%) were down-regulated in treatments of hexadecane compared to acetate (Figure S8). However, among the up-regulated ones, there are more sRNAs showing large expression fold changes (|Log_2_FC| >3) (20.5%) than that (3.3%) in the down-regulated ones (Figure S8), indicating more intense expression response of sRNAs during the alkane metabolism. Meanwhile, among the top 50 highly expressed lists of each condition, most of the sRNAs (36/50) significantly up-regulated in alkane, but only a few sRNAs (13/50) up-regulated in non-alkane treatments (Figure 6), which further supports the more active responses of sRNAs in alkane utilizing. Interestingly, there was a higher proportion of intesRNAs (12% vs. 4%) among the up-regulated sRNAs than the down-regulated ones in the alkane treatments (Figure S8), suggesting that more integenic sRNAs participate in the alkane metabolism. In line with the above active sRNA responses to alkane, the key sRNA-chaperone Hfq was also up-regulated (∼1.8 times) in alkane (Table S17), which can assist the functioning of sRNAs during alkane utilization.

Among the actively responded sRNAs, there are three core sRNAs, i.e., tmRNA, M1 RNA and 6S RNA (Figure 6), which can be further deduced their roles in alkane metabolic regulations based on their known conservative functions. All the three core sRNAs are among the highly expressed sRNAs, in which tmRNA is the most abundant sRNA in either carbon source (Figure 6 and Table S16). The tmRNA and M1 RNA were significantly up-regulated while the 6S RNA was down-regulated in alkane treatments (Figure 6). Given that the important roles of tmRNA in bacterial ribosome rescue and stress responses (Janssen and Hayes, 2012; Guyomar et al., 2021), and bacterial alkane metabolism is usually associated with the membrane and oxidative stresses (Park et al., 2020), we propose that the up-regulated tmRNA may be indirectly implicated in the global regulation of alkane metabolism through mitigating the stresses during the alkane degradation. The M1 RNA as the key component of RNase P that is a major player in the processing, maturation and decay of post-transcriptional RNA metabolism (Mohanty and Kushner, 2018 and 2022), its up-regulation indicates the actively sRNAs-mediated RNA turnovers during the alkane metabolism. Meanwhile, the 6S RNA is an important global repressor of transcription initiations (Chen et al., 2017; Wassarman, 2018), and thus its down-regulation in alkane means more genes would increase their transcriptions. In line with this inference, more genes regardless of sRNAs or mRNAs are detected to significantly increase their expressions responding to the alkane compared to the control (Tables S16 and S17). In other words, *A. dieselolei* may globally switch the expression of related genes at transcriptional level through the core 6S RNA to adapt the alkane utilization. Besides the 6S RNA riboregulator, other key protein-regulators may also participate in the process of transcriptional expression switching between distinct carbon sources. For instance, the H-NS (histone-like nucleoid structuring protein) DNA-binding protein (B5T_00005), as a major genome-wide transcriptional silencer (Ishihama and Shimada, 2021), was also remarkably down-regulated in alkane (Table S17), suggesting the coordinated regulations of sRNA- and protein-regulators during alkane metabolism.

### 2.10. CsrA-related sRNAs involved in alkane metabolism through diverse mechanisms

By far, the most well-known alkane metabolism regulatory mechanism mediated by sRNA is the multi-tier regulating strategies of Crc/Hfq system in *Pseudomonas* (Moreno and Rojo, 2019). However, there is no homologous protein of Crc in *Alcanivorax*. Another key global regulator CsrA (carbon storage regulator; also named Rsm, repressor of secondary metabolism) is widely distributed in this genus and shows a highly consistent synteny of ‘tRNA-CsrA-Aspartate kinase’ with neighbour genes (Table S12), implying its conserved function within *Alcanivorax*. CsrA mainly inhibit translation initiation through binding a conserved GGA motif in the 5’-UTR of mRNA, and the inhibition can be relieved by CsrA-antagonized sRNAs (e.g., CsrB and CsrC) through molecular mimicry (Potts et al., 2017). As a key global regulator, CsrA has been reported to regulate various physiological processes like carbon metabolism, iron metabolism, motility, cell envelope, secondary metabolism and biofilm formation (Yakhnin et al., 2013; Kulkarni et al., 2014; Altegoer et al., 2016; Potts et al., 2017), and the involvement of CsrA in bacterial alkane metabolism has never been reported before.

With the help of CSRA_TARGET program (Kulkarni et al., 2014), the potential targets of CsrA (B5T_03095) were first predicted in *A. dieselolei* (Table S18). Interestingly, the result indicated that some mRNAs of genes involved in the alkane utilization related pathways might be targeted by CsrA, such as *rubA* (B5T_04349) in the electron transfer during alkane hydroxylation, *aldH* (B5T_00039) and *exaA* (B5T_01640) in the aldehyde and alcohol oxidations, *proB* (B5T_03726) in the proline synthesis which is the key component of the lipopeptide biosurfactant generated by *A. dieselolei* B-5 (Qiao and Shao, 2010), and the genes in the *β*-oxidation, such as *fadB* (B5T_01660) and *acdA* (B5T_04219) (Table S18). In addition, genes involved in iron metabolism, motility, central carbon metabolism and cell envelope were also in the targets list (Table S18), similar as previously reports in model bacteria *E. coli* and *B. subtilis* (Altegoer et al., 2016; Potts et al., 2017). These results suggested that the global protein-regulator CsrA may directly participate in the alkane metabolic pathways.

Importantly, combined the InvenireSRNA prediction (Fakhry et al., 2017), Rfam alignment and secondary structure analysis, we identified two putative <u>Csr</u>A-related s<u>R</u>NAs (named CsrR1 and CsrR2) from the 549 sRNAs of *A. dieselolei* (Figures 4A, 7A to 7C, Tables S10 and S18), which may sequester the CsrA regulator and therefore indirectly involve in the alkane metabolism. CsrR1 originated from the 5’UTR of *hfq* (B5T_00774). Hfq is the most important chaperone interacting with sRNAs in most bacteria (Wagner, 2013; Bar et al., 2021). Our results indicate the intricate links among Hfq, CsrA and CsrA-related sRNA (Figure 7D). However, BLASTN-based analysis showed that homologs of CsrR1 only distribute in *A. dieselolei* and *A. xenomutans* (Figure 4B), implying that CsrR1-mediated regulatory relationships with CsrA and Hfq should be localized.

**Figure 7.**
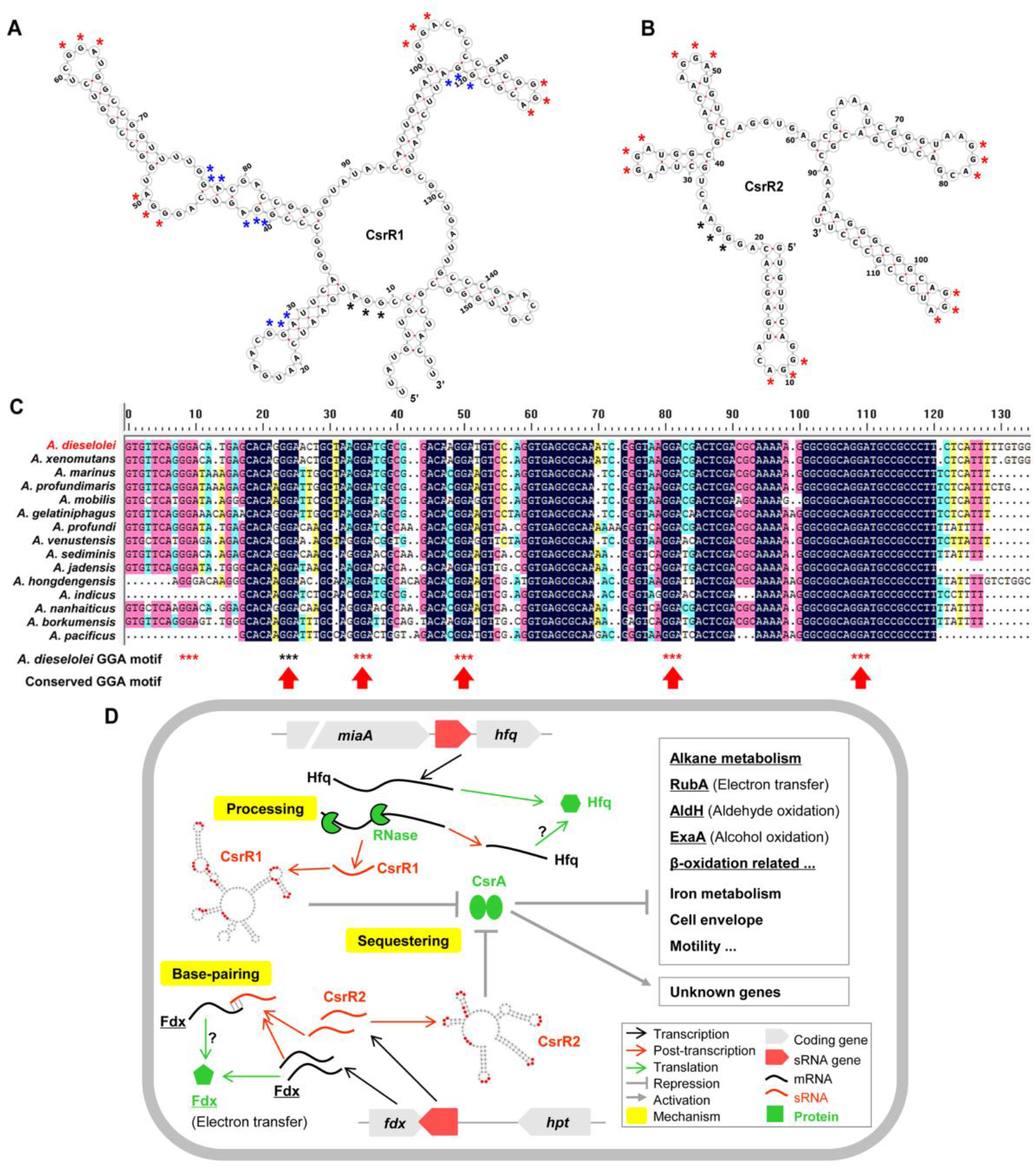
CsrA-related sRNAs and proposed regulation mechanisms in *A. dieselolei*. (**A**) and (**B**) show the secondary structures of CsrR1 and CsrR2 conserved region, respectively. (**C**) The alignments of CsrR2 homologous sequences in different species of Alcanivorax. The potential GGA motifs are marked using asterisks with different colors, red for motifs in loops, black for motifs in single-strand regions and blue for motifs at the junctions of stem-loop. Secondary structures were calculated using RNAfold, and sequences were aligned with DNAMAN (version 8.0). (D) The proposed regulation mechanisms of the two CsrA-related sRNAs in alkane metabolism. The detailed notes are on the bottom right corner. Abbreviations: *miaA*, tRNA dimethylallyltransferase (B5T_00773); *hfq*/Hfq, Host factor (B5T_00774); RubA, Rubredoxin-NAD(+) reductase (B5T_04349); AldH, Aldehyde dehydrogenase (B5T_00039); ExaA, PQQ-dependent dehydrogenase (B5T_01640); *fdx*/Fdx, Ferredoxin (B5T_03135); *hpt*, Hpt domain-containing protein (B5T_03136). The mRNA/protein related with alkane metabolism is underlined. The question mark represents unknown effects.

CsrR2 is independently transcribed through its own promoter recognized by sigma 70 (Table S5), and the homologs are conserved in all *Alcanivorax* species (Figure 4B), indicating a housekeeping and conservative role in this genus. The syntenies of nearby genes are also highly consistent among *Alcanivorax* species (Figure 4C), with a same directional gene encoded Hpt domain-containing protein and a divergent gene *fdx* encoded ferredoxin family protein (Table S12). Intriguingly, CsrR2 is also the antisense RNA of the neighboring *fdx* gene (Figures 4A and 4C), demonstrating that CsrR2 might be a dual-function sRNA through base-pairing and protein sequestering mechanisms (Figure 7D), reminiscent of some previous sRNAs (e.g., McaS and GadY) (Jørgensen et al., 2013 ; Romeo and Babitzke, 2018). Given that the ferredoxin family protein (B5T_03135) is involved in the electron transfer of alkane hydroxylase (Liu et al., 2011), thus CsrR2 may add a regulatory tier to alkane degradation by direct antisense base-pairing with the corresponding mRNA (Figure 7D). Unexpectedly, the same genetic organization of CsrR2 and ferredoxin also occurs in another common alkane-degrader *Pseudomonas*, in which there is a highly conserved orgnization of *fdxA*-*rsmZ*, with *rsmZ* encoding the functional homologue of CsrR2 and *fdxA* encoding a ferredoxin (Lalaouna et al., 2012). The role of CsrA-related sRNAs is highlighted in alkane metabolism.

Based on their different secondary structures and expression patterns, the two CsrA-related sRNAs might play different roles in regulating alkane metabolism. CsrR1 contains five unpaired and four partially paired GGA motifs (Figure 7A), and CsrR2 have six unpaired GGA motifs (Figure 7B). Previous studies have shown that the affinity of CsrA-sRNA interaction is closely related to the secondary structure of sRNA (Duss et al., 2014; Romeo and Babitzke, 2018), and therefore CsrR1 and CsrR2 should have distinct titration abilities to CsrA. Furthermore, the expression of 5’UTR-derived CsrR1 was dependent on its parental mRNA of *hfq*, and both were up-regulated in alkane, while CsrR2 constantly expressed in both conditions (Tables S16 and S17). The distinctive responses to alkane indicate that CsrR2 likely play a fundamental role in antagonizing CsrA, while the up-regulated CsrR1 could indirectly boost translations of the aforementioned alkane metabolism-related genes through further sponging CsrA. Taken together, both CsrR1 and CsrR2 may affect alkane metabolism through diverse mechanisms, which are summarized in Figure 7D.

### 2.11. Candidate sRNAs directly related to the key genes of alkane metabolic pathways

In addition to the above sRNAs that may regulate alkane metabolism globally, more sRNAs that may directly participate in this regulatory process were also analyzed in *A. dieselolei* B-5. To this end, the previously characterized key genes of B-5 (Liu et al., 2011; Li and Shao, 2014; Wang and Shao, 2014), including genes in processes of alkane sensing (*ompS*), chemotaxis (*mcp, cheW, cheR*), transporting (*ompT1-3*), hydroxylation (*alkB1, alkB2, ahpG, almA*), regulation (*almR, cyoD*) and other related processes (*fadB* in *β*-oxidation, *gspE* and *gspF* in putative secretion, *dadA* and *dadB* in haloalkane dehalogenation) were used to determine the alkane metabolism directly related candidate sRNAs according to one of the following three criteria: **i**) intrsRNAs originated from the key genes, **ii**) antisRNAs of the key genes, and **iii**) intrsRNAs or intesRNAs potentially targeting the key genes through *trans*-acting mode (Figure 8A). To reduce the false-positive rate of targets prediction, only the key genes in the top 10 ranking of IntaRNA results are shown here (Figure 8B).

**Figure 8.**
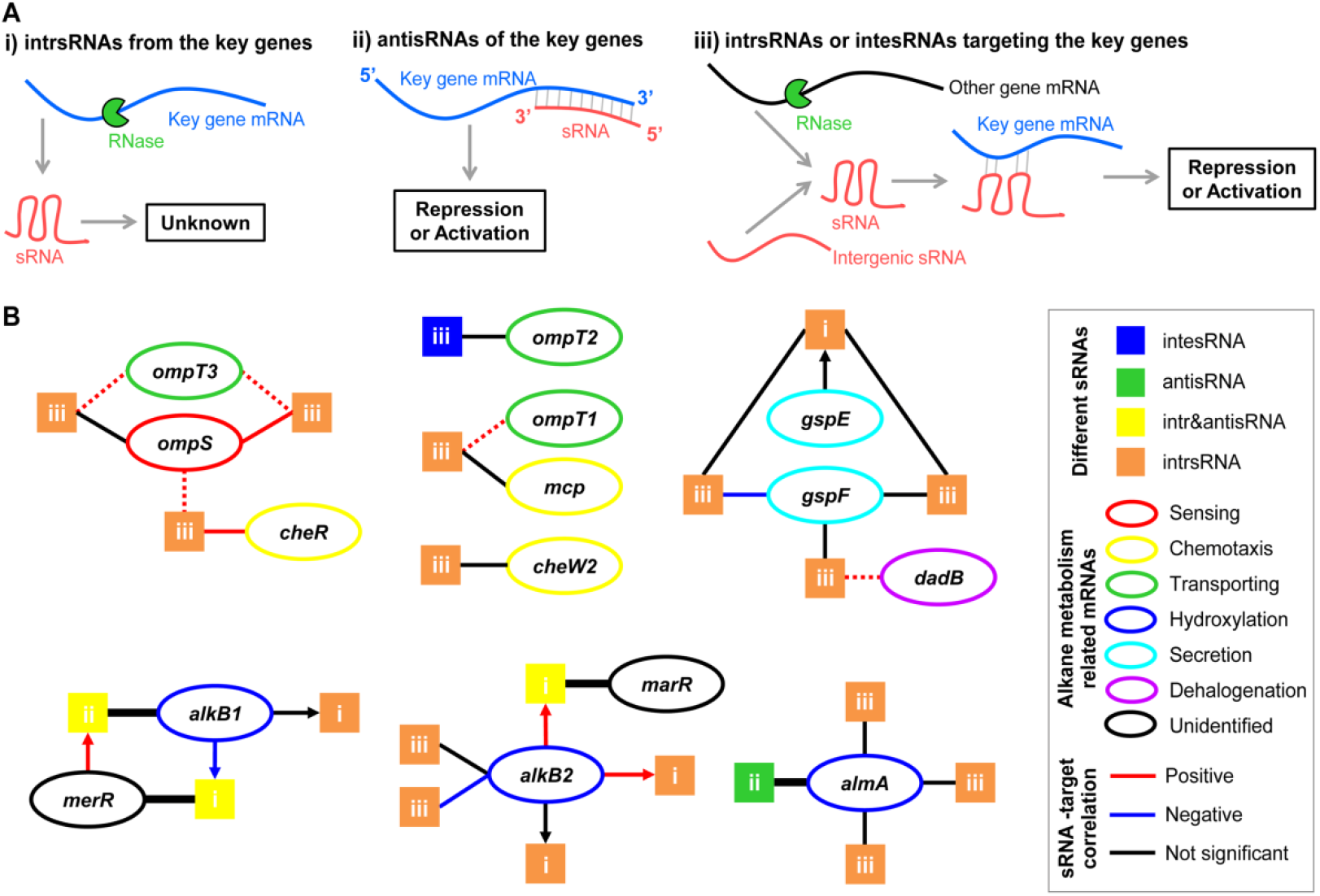
Putative relationships between the key genes in alkane metabolism and directly related sRNAs. (**A**) The three criteria to determine the directly related sRNAs with key genes of alkane metabolism. (**B**) Putative relationships between sRNAs and key genes. Each square represents a unique sRNAs, and the corresponding color shows the location-based classification. The ‘i, ii, iii’ in the squares correspond to the three above criteria. The ovals denote the mRNAs of key genes and are colored according to functions in alkane metabolism, and the related gene names are shown inside. The lines show the matching relationships between sRNA and their targeted mRNAs or sRNAs. The cis-acting relationships were deduced in terms of directly base-pairing for the antisRNA and intr&antisRNA (showing with wider lines), and the trans-acting relationships were inferred by IntaRNA prediction. The key genes in the top 10 most likely targets are shown with solid lines, and dotted lines show the potential targets beyond that range (top 20 to 100). The arrows from mRNAs to sRNAs show the intrsRNAs overlapping with the key genes. The colors of lines or arrows indicate correlationships of the connected both ends based on Spearman correlation analyses of ssRNA-seq expression values in two carbon sources (n=6).

As a result, we identified 21 candidate sRNAs from all the 549 sRNAs, and most of the key genes in various processes of alkane metabolism were targeted by these sRNAs through *cis*- and/or *trans*-acting mechanisms (Figure 8B and Table S19). For instance, OmpS as the essential sensor and chemotaxis trigger of extracellular alkane in *A. dieselolei* (Wang and Shao, 2014), its mRNA is potentially targeted by three *trans*-acting intrsRNAs, and these sRNAs also target other mRNAs of key genes in alkane chemotaxis (*cheR*) and cross-membrane transport (*ompT3*), indicating that multiple processes of alkane metabolism might be closely linked and co-regulated by sRNAs. Moreover, other key genes in alkane transport (*ompT2* and *ompT1*) and chemotaxis (*mcp* and *cheW2*) are also predicted to be directly targeted by intergenic or intragenic *trans*-acting sRNAs (Figure 8B and Table S19). As a powerful alkane-degrader, *A. dieselolei* possesses multiple different alkane hydroxylases to deal with various alkanes (Liu and Shao, 2005; Liu et al., 2011; Wang and Shao, 2014). Three of the identified hydroxylases, including AlkB1, AlkB2, and AlmA for initiating the oxidation of alkanes with different chain lengths and structures, are the parental mRNAs and/or direct targets of several sRNAs (Figure 8B). For the gene *alkB1*, the adjacent gene of MerR family was supposed to be its transcriptional regulator (Liu et al., 2011; Wang and Shao, 2013), here we found two sRNAs were processed from the mRNAs’ 3’-ends of the above two genes and overlapped with the transcripts of *alkB1* and *merR*, indicating mutual *cis*-acting regulations of the two genes at the post-transcriptional level (Figure 8B). Expression of *alkB2* may be negatively regulated by two *trans*-acting intrsRNAs, and *alkB2* is also the parental mRNA of three sRNAs, in which one sRNA from the 3’-UTR may antagonize the neighboring gene expression of a transcriptional regulator of MarR family to indirectly influence the iron metabolism of *A. dieselolei*. AlmA as the key hydroxylase of long-chain alkanes (Wang and Shao, 2014), its expression is intensively regulated by multiple sRNAs through *cis*- and *trans*-acting mechanisms (Figure 8B), reflecting strictly regulations in utilizing long-chain alkanes. In addition, the mRNA of *gspF* which may affect alkane metabolism through secretion shows potential interactions with three *trans*-acting intrsRNAs, and another intrsRNA processed from the 3’-end of *gspE* mRNA may sponge two of the above intrsRNAs (Figure 8B and Table S19), suggesting sRNAs-mediated intricate regulations within an alkane metabolism-related operon (*gspE*-*gspF*). Among all the above 21 candidate sRNAs, there is only one broadly conserved sRNA (sRNA4204314) across the genus *Alcanivorax* (Figure 5 and Table S19), indicating that most of the directly related sRNAs in the alkane metabolic pathways are highly species-specific.

Therefore, sRNA should play a central role in alkane metabolism through diverse and flexible ways. The intricately mixed relationships among sRNA, mRNA and protein, like ‘one-to-many’ or ‘many-to-one’, collectively constitute the competitive alkane metabolic networks in *A. dieselolei*. On the basis of results in this study, the details of sRNA-mediated alkane metabolic regulations still remain to be determined relying on more RNA-RNA/RNA-protein interacting methods in future.

## 3. Materials and methods

### 3.1. Bacterial strain, growth conditions and RNA extraction

The *A. dieselolei* B-5 is a type strain of *Alcanivorax* isolated from the surface water of the Bohai Sea (Liu and Shao, 2005), and the strain is obtained from the Marine Culture Collection of China (MCCC1A00001^T^). It was cultivated in the artificial sea-water medium (ASM) using *n*-alkane hexadecane (0.5%, v/v) and non-alkane acetate (1.0%, w/v) as the sole carbon source, respectively. The mid-log phase (OD_600_ ∼1.0) bacterial cells were harvested at 4°C by centrifugation, then the pellets were lysed in 1 mL of TRIzol Reagent (Invitrogen, USA), and all samples were subsequently stored at -80°C until RNA extraction. Three biological replicates were collected for each condition. The total RNA was extracted using TRIzol reagent according to a previous description (Li et al., 2015). More details are listed in the Supplementary file.

### 3.2. Multiple types of RNA-seq libraries preparation and sequencing

Three types of RNA-seq libraries, including differential RNA-seq (dRNA-seq), fragmented strand-specific RNA-seq (ssRNA-seq) and ribosome profiling sequencing (Ribo-seq), were prepared to target the 5’-ends of transcripts with single-nucleotide resolution, to determine the expression ranges and levels of transcripts, and to evaluate the coding potential of sRNA candidates, respectively.

The dRNA-seq libraries were prepared according to a previous description which could distinguish the primary and processed transcripts through a 5’P-dependent terminator exonuclease (TEX) treatment (Bischler et al. 2015), and a procedure of RNA size (∼50-to 500-nt) selection was added in the process to enrich the small RNAs. After that, the Illumina sequencing was performed on a HiSeq 4000 platform at the Cloud-Seq Biotech (Shanghai, China) according to the manufacturer’s instructions. The dUTP-based strand-specific ssRNA-seq libraries were constructed according to the previous protocols (Parkhomchuk, et al. 2009; Levin et al., 2010) at Majorbio Bio-Pharm Technology (Shanghai, China). Then a HiSeq 4000 platform (Illumina, San Diego, USA) was used for paired-end (2×150 bp) sequencing following the manufacturer’s instructions. The Ribo-seq libraries preparation and sequenceing were performed according to previous studies (Ingolia et al., 2012; Wang et al., 2015) in Gene Denovo Biotechnology (Guangzhou, China). The cDNA libraries were sequenced using the Illumina HiSeq X10 sequencing system. More descriptions on the library construction are detailed in the Supplementary file.

### 3.3. Reads mapping of multiple RNA-seq data

For all the RNA-seq data, the sequencing reads were quality checked according to Illumina standards and then converted into FASTQ format. The READemption tool (version 2.0.1) was used to perform the read mapping with its subcommands of ‘create’ and ‘align’ against the reference genome of *A. dieselolei* B-5 (Lai et al., 2012; Förstner et al., 2014). When aligning with the genome, the parameters of ‘-Q 20’ and ‘-l 20’ were set to exclude the sequences with low sequencing qualities (Phred scores < 20) and shorter than 20-nt, and an extra parameter ‘-c’ was added in the read mapping of dRNA-seq to clip the polyA tails. For dRNA-seq, the single end data targeted the 5’ RNA ends were used for read mapping, while both ends data of ssRNA-seq and Ribo-seq were used for the read mapping.

### 3.4. Visualization of aligned reads

To visualize the mapped results of dRNA-seq and correspongding ssRNA-seq, the READemption subcommand ‘coverage’ was used to generate strand specific coverage files based on the read alignments with default parameters. Finally, the data were visualized in the Integrative Genomics Viewer (IGV, version 2.3.68) (Thorvaldsdóttir et al., 2013), with the whole genome sequence (fasta format), the genomic annotation file (gff format) and the coverage files (wiggle format, normalized by the total number of aligned reads and multiplied by one million) as inputs.

### 3.5. Identification of TSSs, PSSs and transcripts

The ANNOgesic pipeline (version 1.1.4) was used to predict and annotate numerous features through integrating a suite of tools based on the dRNA-seq and ssRNA-seq data (Yu et al., 2018), and all parameters were kept at the default values if not specified. The TSSs and PSSs were identified using the integrated TSSpredator tool in ANNOgesic (subcommand: ‘annogesic tss_ps’) with the programs of ‘TSS’ and ‘PS’ respectively, in which the relative enrichment of reads was compared between the TEX-treated and TEX-untreated samples (Yu et al., 2018; Ryan et al., 2020). The identified TSSs and PSSs were further improved by manual curation based on the visualized read coverage plots in IGV. The coverage-based transcript detection strategy in ANNOgesic (subcommand: ‘annogesic transcript’) was used to predict the transcripts through the input of ssRNA-seq data.

### 3.6. Determination of sRNA candidates

To detect the sRNA candidates, we first used the ANNOgesic’s subcommand ‘annogesic srna’ to scan all the possible sRNAs (both intergenic and CDS-overlapped sRNAs) though adding a parameter of ‘--detect_srna_in_cds’ with the inputs of the transcripts, coverage files, TSSs, PSSs and genome annotation. Meanwhile, the promoter table and terminator file generated by the subcommands of ‘annogesic promoter’ and ‘annogesic terminator’, together with the parameters of ‘--compute_sec_structures’ and ‘--srna_database_path’ were also included in the analyzing process to assit the filter of sRNAs. After that, each of the obtained sRNA candidates was manually verified based on the distributions, abundances and peak shapes of mapped reads in the IGV visualization.

In addition, to overcome the potential bias of the ANNOgesic results due to the selection of parameters, we also manually inspected the distribution of all the mapped reads in IGV across the whole genome to complement the missed sRNA candidates. As a result, only the sRNA candidates that fit all the following criteria are included in the final sRNAs list: **i)** sRNAs have mapped reads both in the dRNA-seq and ssRNA-seq results (to exclude the false-positive introduced by single strategy of library construction); **ii)** the sRNAs’ abundances normalized by total reads number in ssRNA-seq are ≥ 10 at least in one carbon source condition (to exclude the very low abundance); and **iii)** the peak plots of sRNAs are with good shapes, i.e., there are TSSs or PSSs at the 5’-ends, and the 3’-ends are with terminators, PSSs or sharp coverage decreases (to lower the interference from degrading fragments and overlapped transcripts) (Ryan et al., 2020; Ponath et al., 2021).

### 3.7. Analyzing of sRNA primary sequences and secondary structures

The CD-HIT was used to cluster the primary sequence similarity of sRNAs, with thresholds of over 70% sequence identity and 90% sequence length (parameters: -c 0.7, -aS 0.9 and default for others) (Fu et al., 2012). The lengths and G+C contents of sRNAs, CDSs, rRNAs and tRNAs were calculated by BioEdit (version 7.2.5) according to their sequences (Hall et al., 2011). The minimum free energy (MFE) of RNAs was calculated using RNAfold integrated in the RNA Workbench 2.0 with the default settings (Galaxy Version 2.2.10.4, https://rna.usegalaxy.eu/) (Lorenz et al., 2011; Fallmann et al., 2019). To remove the bias from the sequence length, the normalized MFE (NMFE) was calculated by dividing the RNA sizes (Barik and Das, 2018; Dar and Sorek, 2018). PlotsOfData was used to draw boxplots of the above features (Postma and Goedhart, 2019), and their distributions were compared using a Wilcoxon rank sum test calculated by ‘PMCMRplus’ package in R. The RNAfold web server (http://rna.tbi.univie.ac.at//cgi-bin/RNAWebSuite/RNAfold.cgi) was used to predict and visualize the centroid secondary structures of sRNAs with the default parameters (Lorenz et al., 2011).

### 3.8. Motif analyzing of the sequences neighboring TSSs and PSSs

The putative promoter motifs were detected by scanning the 50-nt upstream of TSSs using MEME (https://meme-suite.org/meme/tools/meme) (Bailey et al., 2015; Yu et al., 2018). The upstream 100-nt sequences of TSSs were further analyzed by the iPromoter-2L (Liu et al., 2018), which can provide a high predictive power for different types of bacterial promoters (Cassiano and Silva-Rocha, 2020). The consensens motif of the sequences spanned up- and down-stream 10-nt of the PSSs was aligned and visualized with WebLogo 3 (http://weblogo.threeplusone.com). The above sequence extraction is carried out using a customed perl script.

### 3.9. Coding potential evaluation of sRNAs

GeneMarkS-2 was used to evaluate the possible coding potential of sRNAs by the default parameters (Lomsadze et al., 2018), and the predicted coding sequences were submited to Pfam for homology searching (http://pfam.xfam.org/). The filtered high quality reads of Ribo-seq were mapped to the detected sRNAs using Bowtie2 (Langmead and Salzberg, 2012). A custom ORFfinder search was conducted in the mapped sRNAs, and only ORFs with lengths ≥20, with complete start (ATG) and stop codons, were retained as potential sORFs. Blastp was used to homology searches of the potential sORFs against the non-redundant protein sequences (nr) (Johnson et al., 2008).

### 3.10. sRNAs annotation and homologs analysis

All the sRNA candidates were aligned and annotated within the Rfam database (version 14.8, https://rfam.org/search/batch). The 15 reference genomes of *Alcanivorax* were downloaded from NCBI Datasets (https://www.ncbi.nlm.nih.gov/datasets/genomes). Then BLASTN was used for similarity searches of the Rfam-annotated sRNAs against the reference genomes with relative loose parameters (−perc_identity 50 -qcov_hsp_perc 50 -evalue 0.00001) for the identity and coverage (Altschul et al., 1997). The gene synteny and Rfam annotation of the obtained sRNA homologs were performed to further verify their conservation. For the gene synteny analysis, the locations of sRNA neighboring genes are determined by the BLASTN result of hsp (high-scoring segment pair) hit regions, and the gene annotations are based on the Prokaryotic Genome Annotation Pipeline (PGAP) (Tatusova et al., 2016). To reduce the false-positives, except the Rfam-annotated sRNAs, we adopted more stringent BLASTN parameters (−perc_identity 75 -qcov_hsp_perc 75 -evalue 0.00001) to search the homologs of the remaining sRNAs in *Alcanivorax* according to a previously described threshold (Ponath et al., 2021).

### 3.11. Expression analysis of sRNAs

We used the ‘gene_quanti’ and ‘deseq’ subcommands of READemption to calculate the reads number overlapping the identified sRNAs and the differential expression in alkane and non-alkane conditions with the default parameters (Förstner et al., 2014; Love et al., 2014). The expression level and change are represented by TPM (Transcripts per Million) normalized read counts and Log2FoldChange with adjusted P value. Differential expressed genes were defined as the adjusted P value < 0.05 and the absolute values of TPM Log2FoldChange ≥1 between the two conditions. The heatmap and volcano plots were used to visualize the gene expression levels and changes via the Heatmapper and VolcaNoseR (Babicki et al., 2016; Goedhart and Luijsterburg, 2020), respectively.

### 3.12. Predictions of CsrA-related sRNAs and CsrA targets

CsrA-related sRNAs were identified with the assistance of the R package “InvenireSRNA”, which integrated sequence- and structure-based features to train machine-learning models for detecting the bacterial sRNAs in the CsrA pathway (Fakhry et al., 2017). All the sRNA candidates and the extracted intergenic sequences of the whole genome were scanned using the InvenireSRNA to discover all the possible CsrA-related sRNAs in *A. dieselolei*. The putative CsrA-repressed targets across the whole genome were identified using the predictive algorithm CSRA_TARGET (Kulkarni et al., 2014), which is based on sequence feature of the conservative CsrA binding motif (ANGGA) in regions around the Shine-Dalgarno (SD) sequence.

### 3.13. Targets prediction of sRNAs

The IntaRNA 2.0 program, which enables fast and accurate prediction of RNA-RNA interactions by integrating seed constraints and interaction site accessibility (Mann et al., 2017), was used to predict the mRNA targets of the *trans*-acting sRNAs in B-5 genome. With the inputs of sRNA sequences and B-5 genomic information (RefSeq ID: NC_018691), the ‘whole genome mode’ of IntaRNA was selected to target the potential regions of annotated genes using the default parameters (http://rna.informatik.uni-freiburg.de/IntaRNA/Input.jsp).

## 4. Conclusions

Here, combined the advantages of dRNA-seq and RNA size selection strategy, we comprehensively captured the high-resolution sRNAs landscape in a marine alkane-degrading bacterium *A. dieselolei* B-5. These sRNAs can originate from nearly everywhere of the genome through ways of promoter-driven transcription or post-transcriptional RNA processing. RNase E likely plays a key role in the processing and maturation of sRNAs. Most of the sRNAs have no sORF-coding potential and are species-specific distribution in *Alcanivorax*. Some core sRNAs, including 6S RNA, M1 RNA and tmRNA, are first revealed responding to alkane, which likely reprogram the global gene expressions at multiple levels of transcription, post-transcription, and translation to benefit the alkane utilization. Two novel CsrA-related sRNAs may also complementarily regulate alkane metabolism through sponging the global translation repressor CsrA and in concert with other mechanisms. Species-specific sRNA-mediated regulations may be widely involved in key processes of alkane sensing, chemotaxis, transporting, and hydroxylation. Altogether, both core and local/specific sRNAs collaboratively reshape the gene expressions at various levels through diverse mechanisms to efficiently optimize the alkane utilization in B-5 (Summarized in Figure 9). These results indicate that sRNAs are at the hubs of regulations, not only fast respond to the change of carbon sources globally via core sRNAs, but also precisely target and coordinate the key pathways in alkane metabolism by multiple specific sRNAs. Our study opens up an important avenue for exploring the sRNA-mediated regulatory networks and will stimulate further work in identification of new functional sRNAs and novel regulatory mechanisms in alkane metabolism.

**Figure 9.**
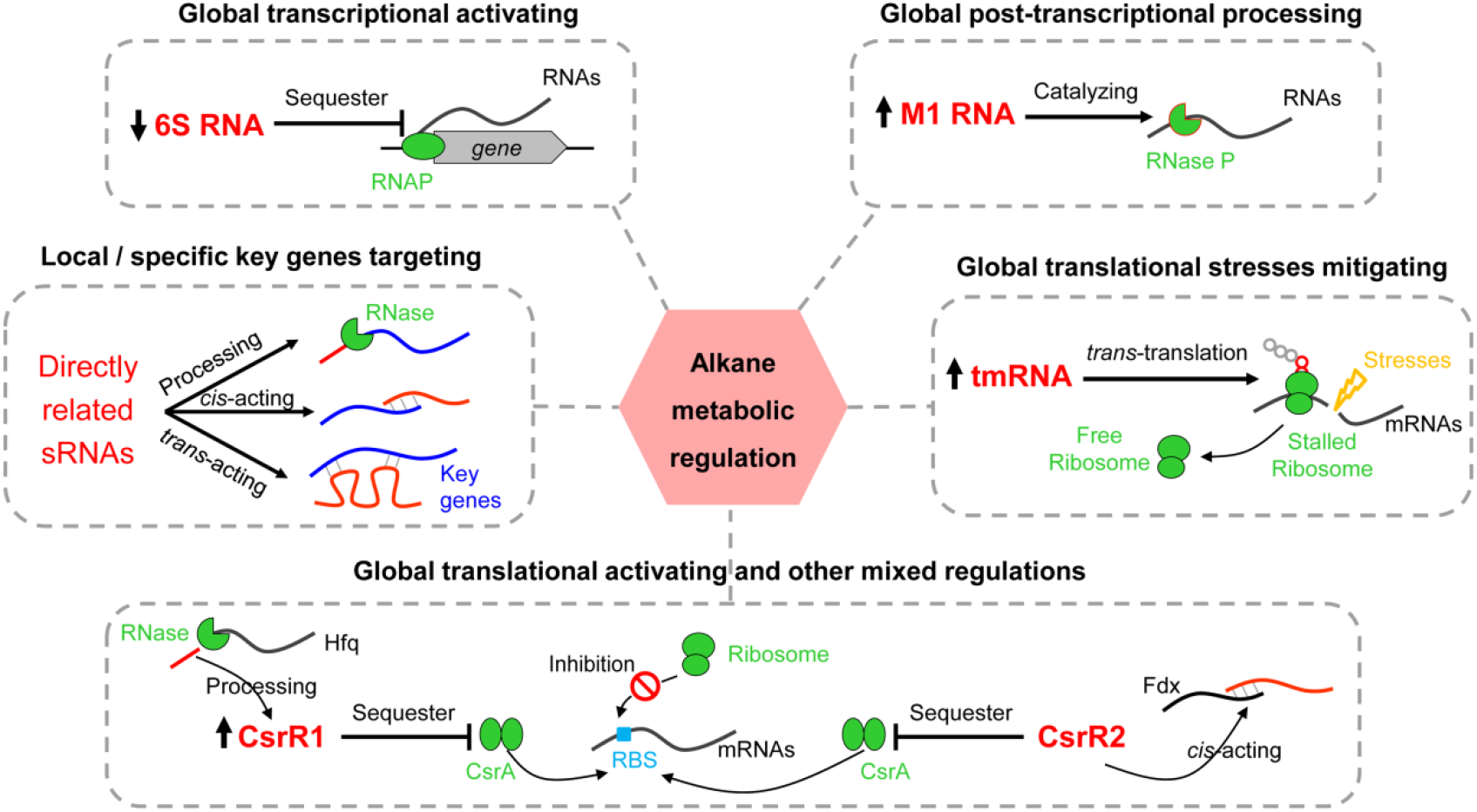
Proposed sRNA-mediated alkane metabolic regulations in *A. dieselolei* B-5. The key related sRNAs (red and largest font) and their potential regulating mechanisms are shown in each dotted box. The significant expression changed sRNAs responding to the alkane are marked with up and down arrows before them, representing for up- and down-regulated expressions, respectively. The possible regulating effects of the sRNAs are indicated above each dotted box (black and bold font). Abbreviations: RNAP, RNA polymerase; Hfq, Host factor; Fdx, Ferredoxin; RBS, ribosome binding site.

## Author Contributions

Conceptualization, G. W. and Z. S.; investigation, G. W., S. L. and Z. W.; supervision, Z. S., J. H. and W. W.; formal analysis, G. W., S. Y. and K. Z.; visualization, G. W.; writing-original draft, G. W.; writing-review and editing, G. W., Z. S., S. Y. and W. W.; funding acquisition, Z. S. and W.W. All authors have read and agreed to the published version of the manuscript.

## Data Availability Statement

The rawdata have been deposited in the Genome Sequence Archive (Chen et al., 2021) in the National Genomics Data Center (NGDC, https://ngdc.cncb.ac.cn/gsa) under the accession numbers of CRA008650 (ssRNA-seq), CRA008652 (dRNA-seq) and CRA008653 (Ribo-seq), respectively. Other available data are uploaded to the figshare, including the data of IGV visualization, the ANNOgesic related outputs and the results of IntaRNA-based targets prediction with the links of https://doi.org/10.6084/m9.figshare.18094298, https://doi.org/10.6084/m9.figshare.21159580 and https://doi.org/10.6084/m9.figshare.21195049, respectively.

## Acknowledgements

We thank Sung-Huan Yu and Jiang Liu for their help during the bioinformatic analyses.

