## Supplementary file for "High-resolution small RNAs landscape provides insights into alkane adaptation in the marine alkane-degrader *Alcanivorax dieselolei* B-5"

**Supplementary Materials and Methods**

**Bacterial strain cultivation and** **RNA extraction**

*Cultivation.* The ASM was prepared as described previously (Liu and Shao, 2005), amended with lower pH (7.2) and less MgSO_4_·7H_2_O (3.5 g/L) to avoid medium precipitation. All cultures were incubated at 28^o^C and spun at 180 r.p.m. for growth growth monitoring.

*RNA extraction*. Briefly, cell pellets corresponding to a total amount of ~3.0 OD_600_ were lysed using 1 mL of TRIzol, followed by adding 200 μL of chloroform and shaking vigorously. After centrifugation at 4^o^C, 12,000 g for 15 min, the supernatant was fully mixed with 500 μL of isopropanol. After another round of centrifugation, the pelle was washed twice using 70% ethanol, then air-dried and dissolved in RNase-free water. RNA purity and integrity were assessed using a Nanodrop 2000 Spectrophotometer (Thermo Scientific) and the Agilent Bioanalyzer 2100 system (Agilent Technologies), respectively.

**Multiple RNA-seq libraries preparation and sequencing**

*dRNA-seq*. Briefly, the residual genomic DNA and ribosomal RNAs were first removed by DNase I (NEB) and Ribo-Zero^TM^ rRNA Removal Kit (Bacteria) (Epicentre) according to the manufacturers’ instructions. Then, the RNA samples were poly(A)-tailed using *E. coli* poly(A) polymerase (NEB), and terminator^TM^ 5’-phosphate-dependent exonuclease (TEX) (Epicentre) was used to degrade the processed transcripts with 5’-P structure and enrich the primary transcripts with 5’-PPP as previously described (Bischler et al. 2015). For each carbon source condition, two treatments, with TEX (TEX+) and without TEX (TEX-), were carried out after poly(A)-tailing. Finally, all samples were treated with TAP (tobacco acid pyrophosphatase, Epicentre) to transform the 5’-PPP to 5’-P, then the Illumina sequencing RNA adapter was ligated to the 5’-P of TAP-treated RNAs using T4 RNA ligase (NEB). To obtain majority of the sRNAs with accurate 5’-ends, RNAs within 50-500 nt were size-selected using 8% polyacrylamide gel electrophoresis, and no fragmented step was performed before the cDNA library preparation (Bischler et al. 2015; Leonard et al., 2019). The cDNA libraries were constructed as previously described (Bischler et al. 2015), and then the Illumina sequencing was performed on a HiSeq 4000 platform at the Cloud-Seq Biotech (Shanghai, China) according to the manufacturer’s instructions.

*ssRNA-seq*. Briefly, after removal of the residual genomic DNA and ribosomal RNAs as described for the dRNA-seq library, dUTP-based strand-specific libraries were created using the TruSeq^TM^ Stranded Total RNA Library Prep Kit (Illumina) according to the manufacturer’s protocols (Parkhomchuk, et al. 2009; Levin et al., 2010).

*Ribo-seq*. Briefly, chloramphenicol with a final concentration of 100 μg/mL was used for ribosome stalling, and cells were collected by centrifugation and immediately flash frozen in liquid nitrogen together with 400 μL bacterial lysis buffer as described previously (Wang et al., 2015). Then the ribosome footprints (RFs) were recovered according to previous studies (Ingolia et al., 2012; Wang et al., 2015). After obtaining the RFs, the Ribo-seq cDNA libraries were constructed using the NEBNext^®^ Multiple Small RNA Library Prep Set for Illumina^®^ (New England Biolabs, MA, USA) according to the manufacturer’s instructions.

**Experimental objectives and sampling strategies in this study**

| **Objective** | **Library type** | **Sampling strategy** | **Culture condition** |
| --- | --- | --- | --- |
| To accurately identify sRNAs with different origins and locations | dRNA-seq+ ssRNA-seq | Three independent biological replicates with similar OD_600_ values of the mid-log phase were merged into one mixture after RNA extractions for each condition | *n*-hexadecane vs. acetate as the only carbon source |
| To determine the expression boundaries and levels of sRNAs | ssRNA-seq | Three independent biological replicates with similar OD_600_ values of the mid-log phase for each condition | *n*-hexadecane vs. acetate as the only carbon source |
| To evaluate the coding potential of sRNAs | Ribo-seq | Two independent biological replicates with similar OD_600_ values of the mid-log phase for each condition | *n*-hexadecane vs. acetate as the only carbon source |

**Reference genomes information of the 15 *Alcanivorax* species used in this study**

| **Species**  **name** | **Genome**  **size (Mb)** | **G+C**  **content (%)** | **Gene**  **number** | **Genome**  **level** | **Rference** |
| --- | --- | --- | --- | --- | --- |
| *A. dieselolei B-5* | 4.928 | 61.5 | 4468 | Complete | Lai et al. 2012, doi: 10.1128/JB.01813-12. |
| *A. xenomutans JC109* | 4.35 | 61.5 | 3962 | Scaffold | https://www.ncbi.nlm.nih.gov/data-hub/genome/GCF_900217905.1/ |
| *A. marinus SY10-13* | 4.175 | 65 | 3883 | Contig | https://www.ncbi.nlm.nih.gov/data-hub/genome/GCF_016785095.1/ |
| *A. gelatiniphagus MEBiC 08158* | 4.215 | 65.2 | 3887 | Contig | https://www.ncbi.nlm.nih.gov/data-hub/genome/GCF_005938655.1/ |
| *A. profundimaris ST75FaO-1* | 4.036 | 66 | 3777 | Scaffold | https://www.ncbi.nlm.nih.gov/data-hub/genome/GCF_015265435.1/ |
| *A. mobilis MT13131* | 4.1 | 63 | 3782 | Contig | https://www.ncbi.nlm.nih.gov/data-hub/genome/GCF_002864685.1/ |
| *A. venustensis ISO4* | 3.544 | 64.5 | 3393 | Contig | https://www.ncbi.nlm.nih.gov/data-hub/genome/GCF_015356855.1/ |
| *A. jadensis T9* | 3.629 | 58 | 3327 | Contig | https://www.ncbi.nlm.nih.gov/data-hub/genome/GCF_000756655.1/ |
| *A. sediminis PA15-N-34* | 3.798 | 57 | 3451 | Contig | https://www.ncbi.nlm.nih.gov/data-hub/genome/GCF_009601165.1/ |
| *A. nanhaiticus 19-m-6* | 4.133 | 56 | 3806 | Contig | https://www.ncbi.nlm.nih.gov/data-hub/genome/GCF_000756665.1/ |
| *A. profundi MTEO17* | 3.737 | 57 | 3506 | Contig | https://www.ncbi.nlm.nih.gov/data-hub/genome/GCF_003597125.1/ |
| *A. hongdengensis A-11-3* | 3.665 | 60.5 | 3498 | Contig | Lai et al. 2012, doi: 10.1128/JB.01849-12. |
| *A. borkumensis SK2* | 3.12 | 54.5 | 2826 | Complete | Schneiker et al. 2006, doi: 10.1038/nbt1232. |
| *A. indicus SW127* | 3.445 | 62.5 | 3161 | Contig | https://www.ncbi.nlm.nih.gov/data-hub/genome/GCF_003259185.1/ |
| *A. pacificus W11-5* | 4.168 | 62.5 | 3785 | Complete | Lai and Shao 2012, doi: 10.1128/JB.01845-12. |

The data were collected from the NCBI Datasets before the July, 2021.
