## Supplementary Figures S1 to S8 for "High-resolution small RNAs landscape provides insights into alkane adaptation in the marine alkane-degrader *Alcanivorax dieselolei* B-5"

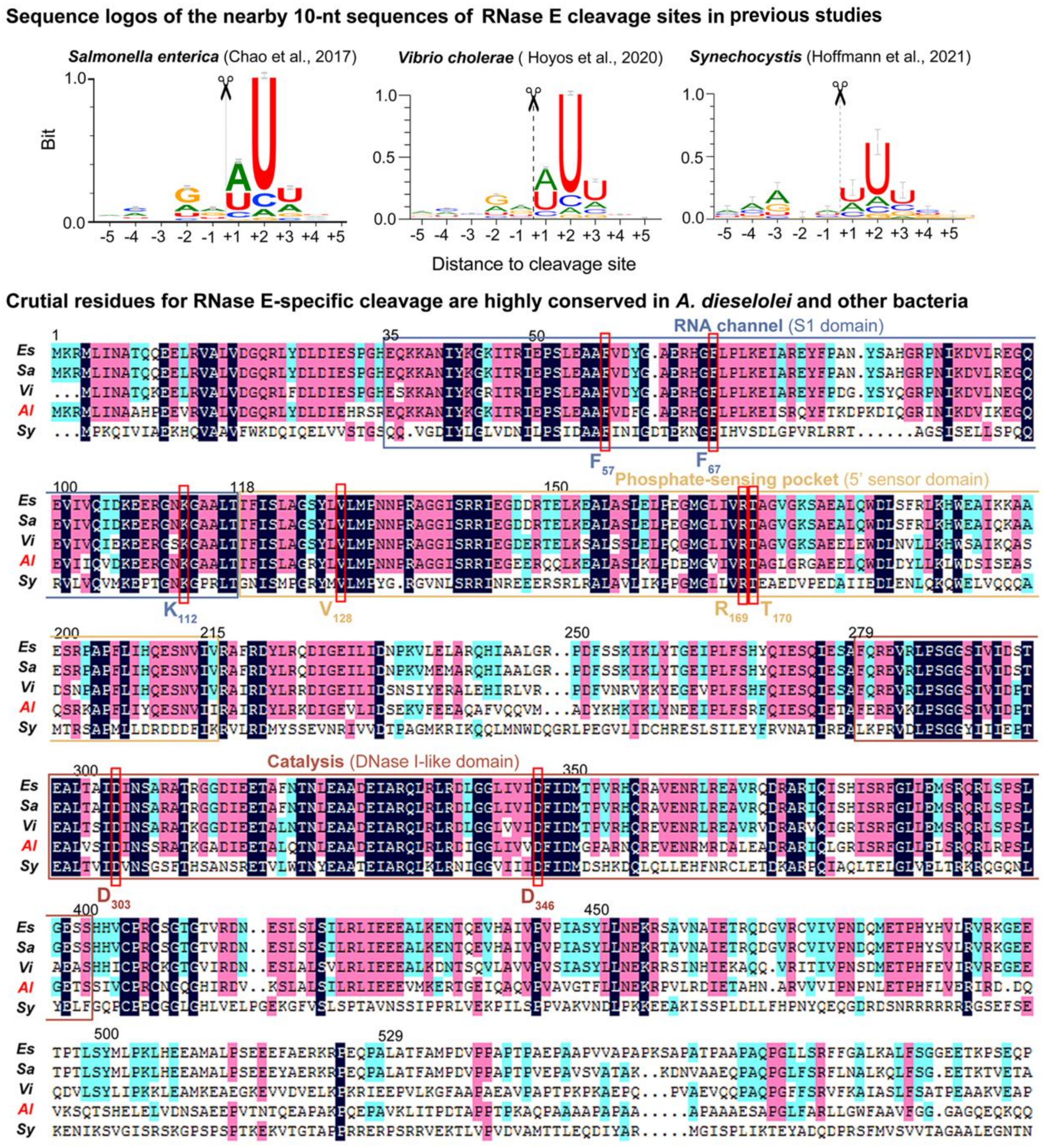


**Figure S1 Sequence logos of the TSSs and PSSs, and key residues in RNase E of different species**

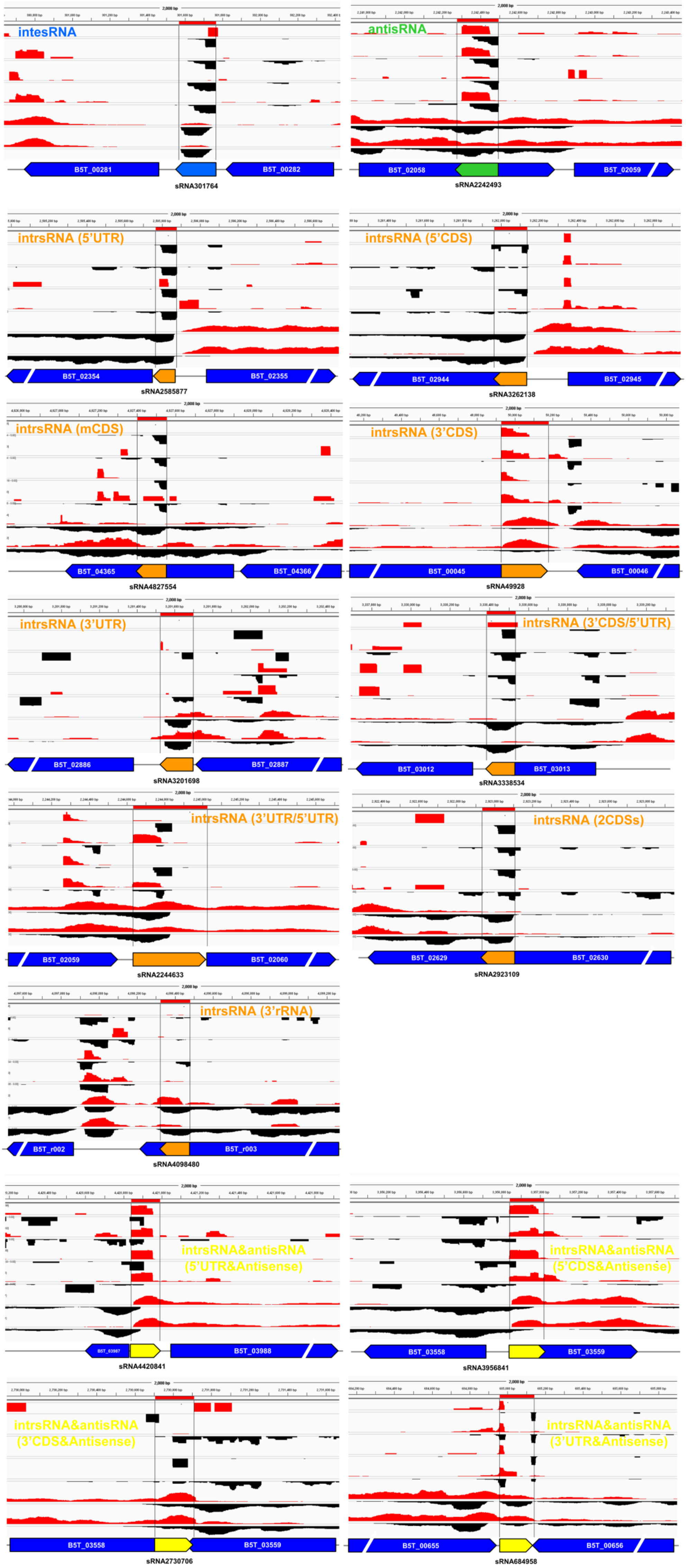


**Figure S2 Examples of peak plots of different sRNA classifications based on the genomic locations**

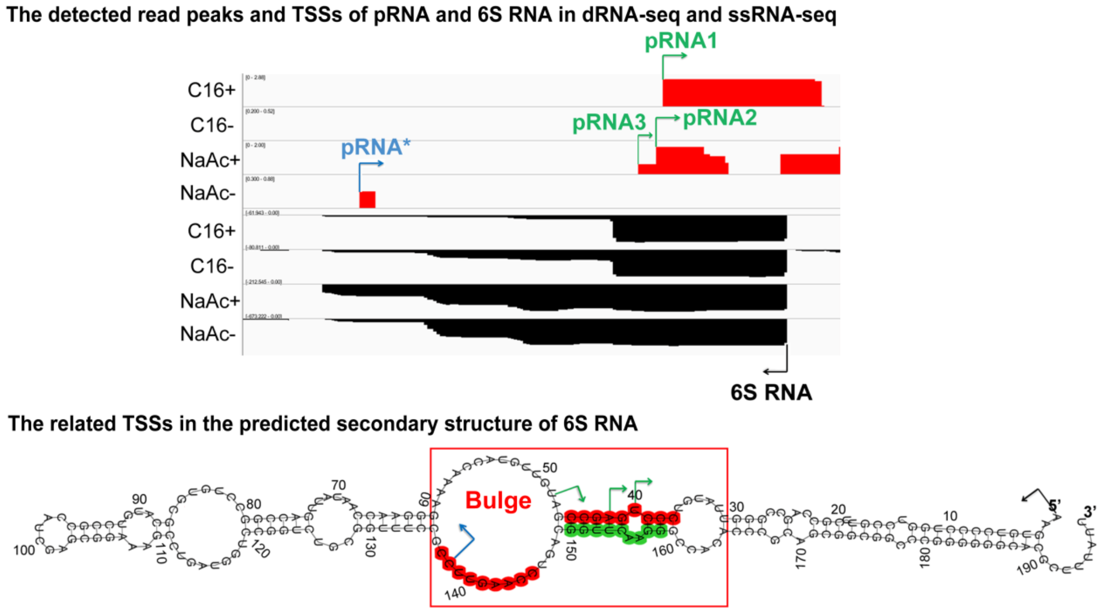


**Figure S3 Putative pRNAs of 6S RNA in *A. dieselolei***

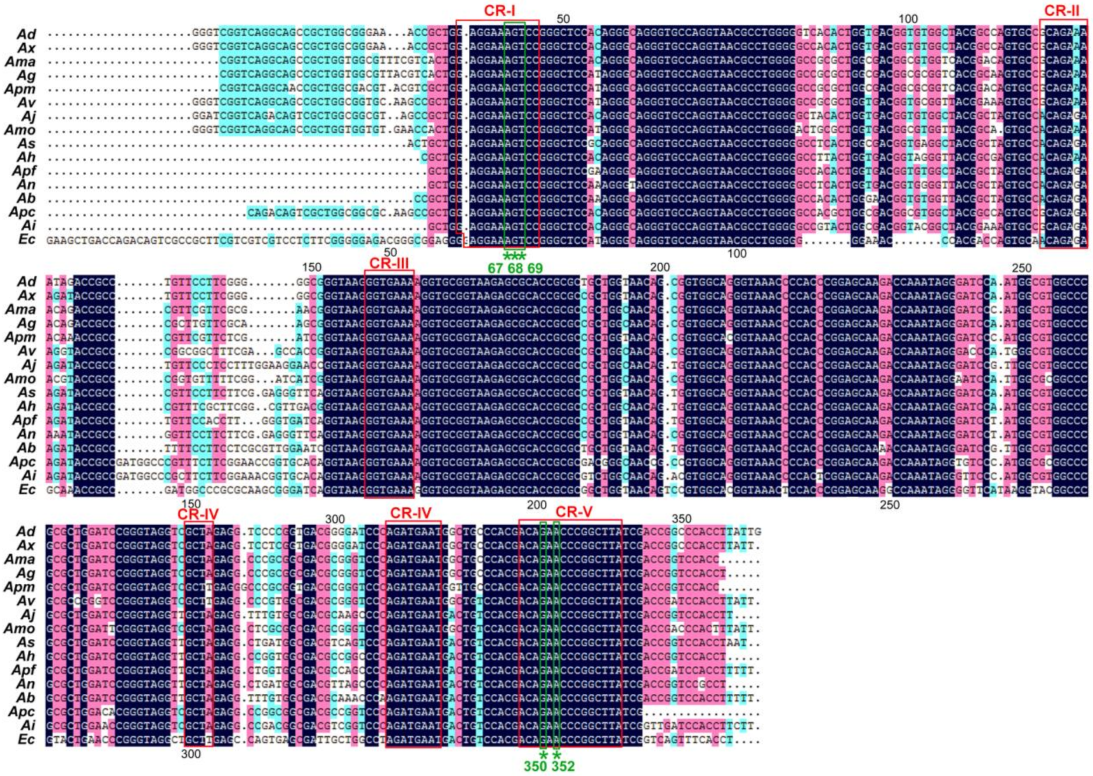


**Figure S4 Conserved regions (CR-I to CR-V) and active sites (*) of M1 RNAs in different *Alcanivorax* species**

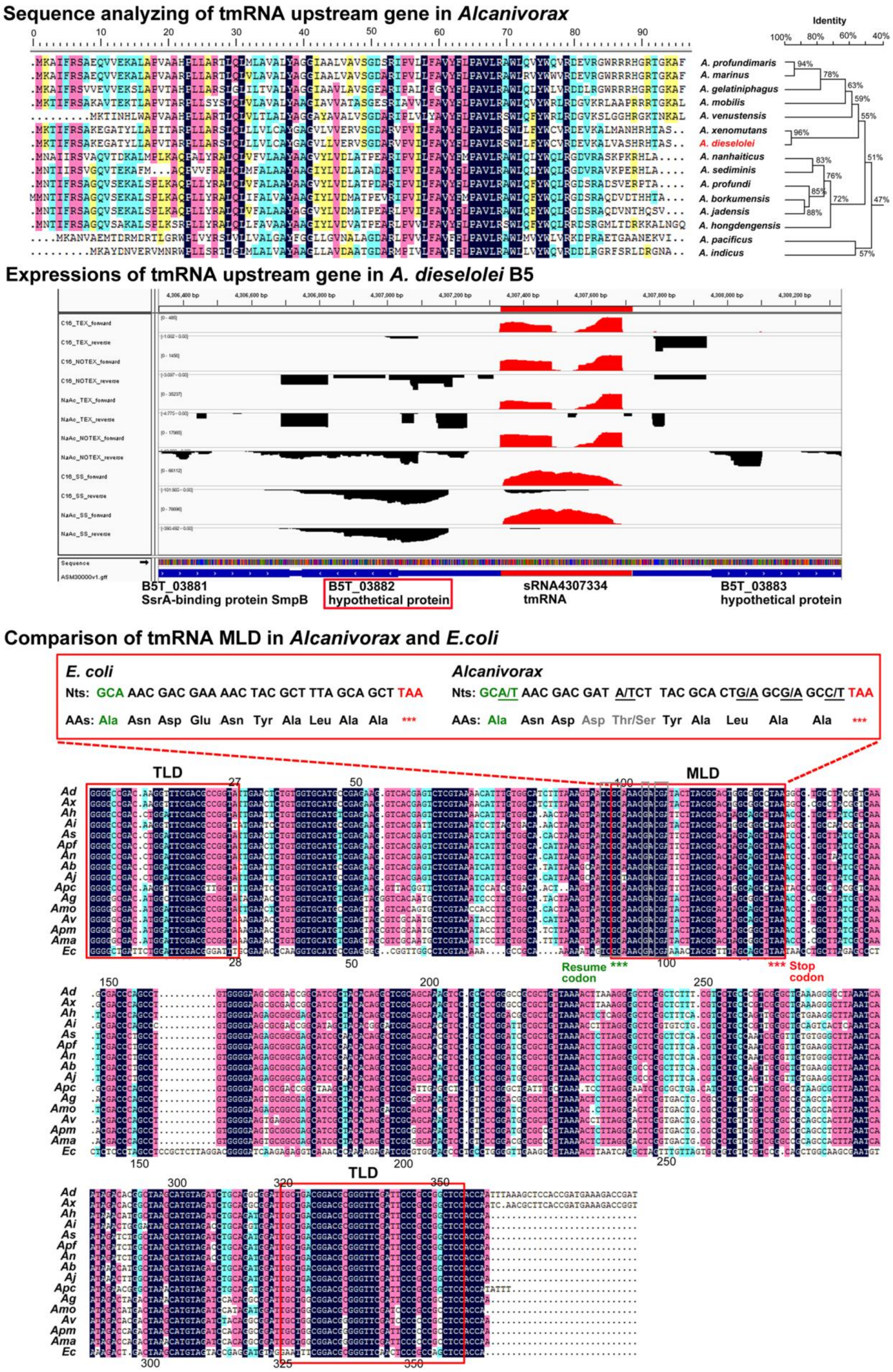


**Figure S5 tmRNA upstream gene and MLD features in different *Alcanivorax* species**

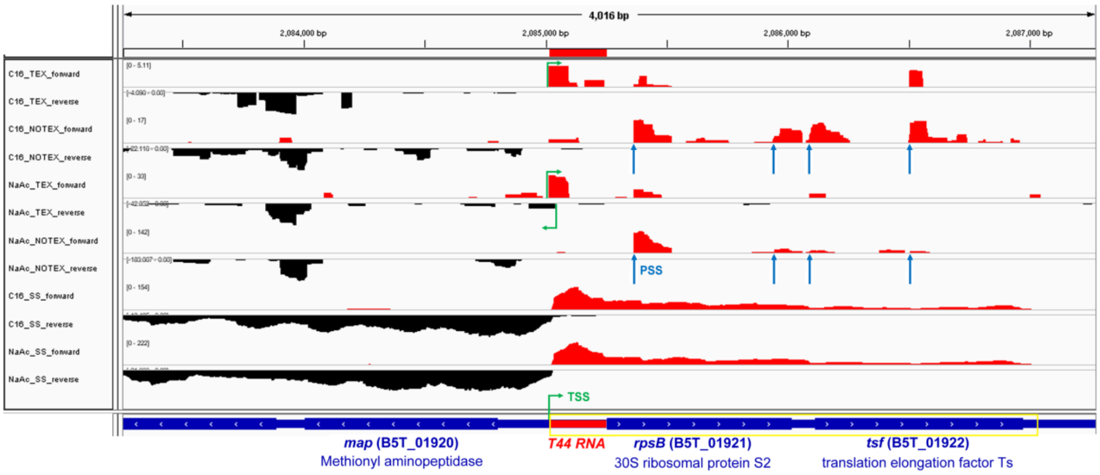


**Figure S6 Expression peaks of T44 RNA and nearby genes**

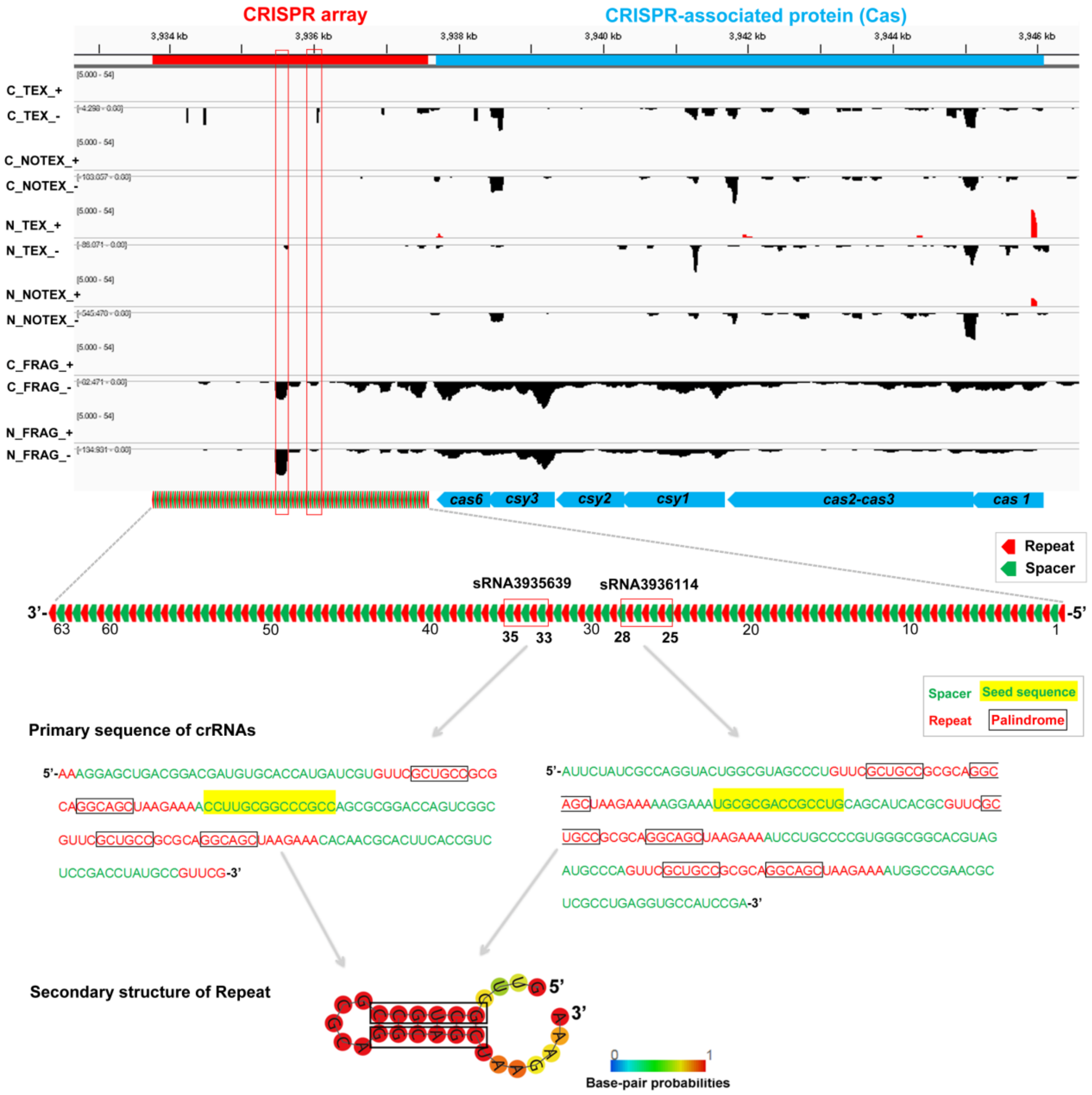


**Figure S7 Expression and identificaion of CRISPR-Cas system in *A. dieselolei***

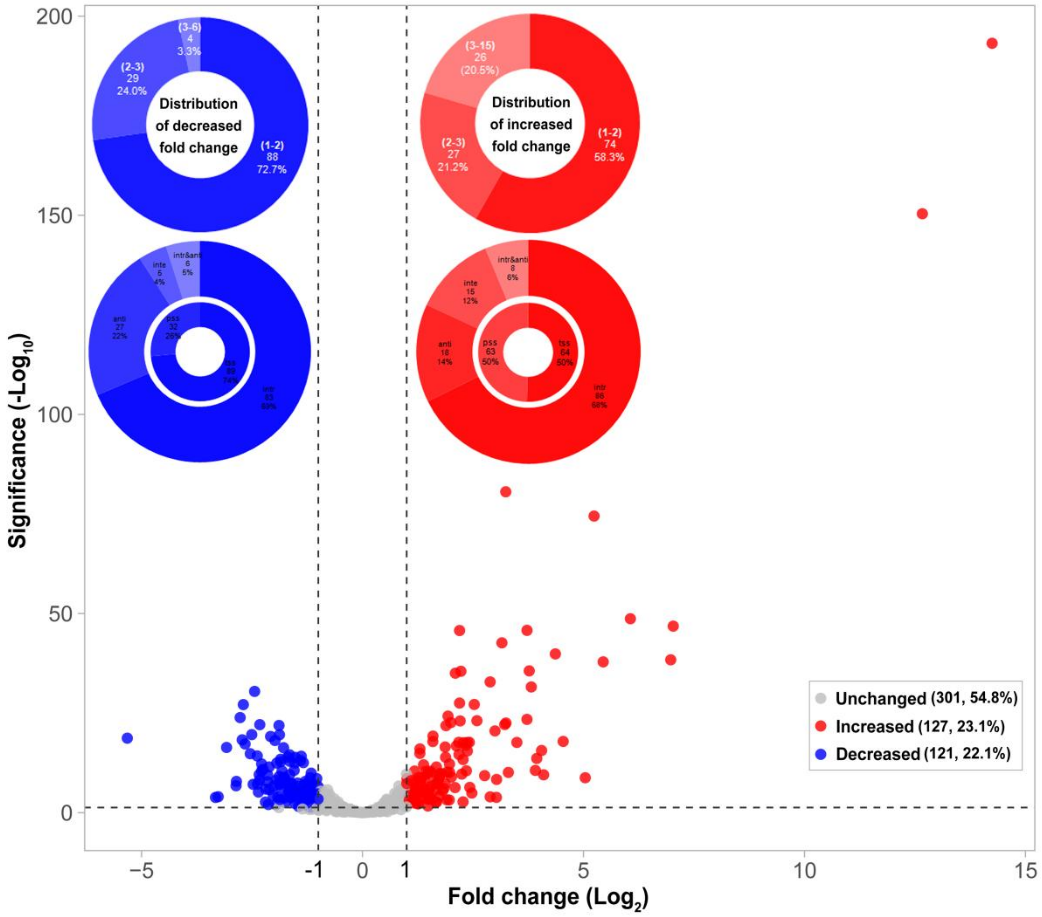


**Figure S8 Volcano plots showing the differential expression in alkane versus acetate conditions**
